# Therapeutic Tumor Macrophage Reprogramming in Breast Cancer Through a Peptide-Drug Conjugate

**DOI:** 10.1101/2024.08.12.607575

**Authors:** Anni Lepland, Elisa Peranzoni, Uku Haljasorg, Eliana K. Asciutto, Maria Crespí-Amer, Lorenzo Modesti, Kalle Kilk, Manuel Lombardia, Gerardo Acosta, Miriam Royo, Pärt Peterson, Ilaria Marigo, Tambet Teesalu, Pablo Scodeller

## Abstract

In triple negative breast cancer (TNBC), pro-tumoral macrophages promote metastasis and suppress the immune response. To target these cells, we engineered a previously identified CD206 (mannose receptor)-binding peptide, mUNO, to enhance its affinity and proteolytic stability. The new rationally designed peptide, MACTIDE, includes a trypsin inhibitor loop, from the Sunflower Trypsin Inhibitor-I. Binding studies to recombinant CD206 revealed a 15-fold lower K_D_ for MACTIDE compared to parental mUNO. Additionally, mass spectrometry showed a 5-fold increase in half-life in tumor lysate for MACTIDE compared to mUNO. Homing studies in TNBC-bearing mice showed that fluorescein (FAM)-MACTIDE precisely targeted CD206^+^ tumor-associated macrophages (TAMs) upon intravenous, intraperitoneal and even oral administration, with no significant accumulation in liver. We coupled MACTIDE to the FDA-approved drug Verteporfin, an established photosensitizer for photodynamic therapy and inhibitor of the YAP/TAZ pathway, to generate a conjugate here referred to as MACTIDE-V. In the orthotopic 4T1 TNBC mouse model, non-irradiated MACTIDE-V-treated mice unexpectedly showed a similar anti-tumoral effect and fewer signs of toxicity as irradiated MACTIDE-V-treated mice, leading to subsequent studies on the laser-independent activity of this conjugate. *In vitro* studies using bone-marrow derived mouse macrophages showed that MACTIDE-V excluded YAP from the nucleus, increased the phagocytic activity and upregulated several genes associated with cytotoxic anti-tumoral macrophages. In mouse models of TNBC, MACTIDE-V slowed primary tumor growth, suppressed lung metastases, increased markers of phagocytosis and antigen presentation in TAMs and monocytes, increasing the tumor infiltration of several lymphocyte subsets. We therefore propose MACTIDE-V as a useful peptide-drug conjugate to modulate macrophage function in the context of breast tumor immunotherapy.

## INTRODUCTION

Established solid tumors favor immunosuppressive and angiogenic phenotypes of macrophages that induce progression and metastasis. Additionally, by secreting cytokines that attract and skew macrophage functions, most tumors can expand the pro-tumoral TAM population, making it the most prominent immune cell type of the tumor microenvironment. Pro-tumoral TAMs are important in TNBC, where they execute several tumorigenic functions^1^ and where antibody-dependent cellular cytotoxicity-based therapies are currently not available due to lack of specific cancer cell receptors. Tackling the tumor-promoting microenvironment through targeting TAMs is therefore an emerging alternative.

To target TAMs, we previously developed the mUNO targeting peptide^2,3^ (sequence: CSPGAK), which binds to the mannose receptor CD206, over-expressed in a subset of pro-tumoral TAMs^4^. In the TNBC 4T1 model, we also showed targeting of mUNO to TAMs with almost no accumulation in liver^5^.

Targeting CD206 is also attractive for diagnosis, as a higher number of CD206^+^ cells in the lymph nodes correlates with onset of relapse in some cancers^6^. Most chemically synthesized systems designed to target CD206 utilize mannose as the recognition moiety. However, as the affinity of mannose is on the low millimolar range^7^, it dictates the need for multivalent presentation in order to achieve binding^8^.

Short linear targeting peptides are selective ligands, that can guide therapeutic or imaging cargos to the tumors^9^. However, short unconstrained peptides have high conformational freedom which translates into poor affinity and proteolytic degradation. The relatively low affinity makes them suitable as multivalent ligands^10^, but in order to develop a monovalent peptide-drug conjugate (PDC), a higher affinity and stability of the targeting moiety is desirable.

Here, we engineered mUNO with the aim of enhancing its affinity and proteolytic stability, to be used in a monovalent format. To this end, we exploited the benefits of the Sunflower Trypsin Inhibitor I (SFTI-1), a conformationally constrained plant-derived peptide, composed of two loops separated by a disulfide bond. The largest loop of SFTI-1 (“primary loop”) inhibits the activity of serine proteases including trypsin and pepsin. The other loop (“cyclization loop”) has been modified by other groups to introduce foreign peptides without affecting the enzyme-inhibiting activity or the oral availability^11^. Additionally, variants of SFTI-1 wherein the cyclization loop is opened display similar inhibition constants to trypsin than the original SFTI-1^12,13^.

We report here a new CD206-binding peptide, MACTIDE, with higher affinity and stability than its predecessor that can be delivered even orally. Secondly, we report a PDC, MACTIDE-Verteporfin (MACTIDE-V), that may be used for light-dependent depletion of CD206^+^ macrophages through photodynamic therapy (PDT) or to reprogram TAMs to an anti-tumoral phenotype through the effect of Verteporfin, an inhibitor of the YAP/TAZ pathway. To our knowledge, MACTIDE-V represents the first PDC to skew TAMs towards an anti-tumoral and anti-metastatic phenotype.

## RESULTS

### 1. Design and molecular dynamics of MACTIDE

To design MACTIDE peptide, we inserted mUNO (sequence CSPGAK) in lieu of the cyclization loop of SFTI-1. Then we removed the head-to-tail cyclization to accommodate a fluorophore/drug on the N-terminus after which, with the intention of increasing flexibility between the cyclization loop and the fluorophore/drug, we added glycine G1 (Fig. 1A) as it is the amino acid with the highest flexibility^14^. To assess if our modifications constrained the mUNO motif in MACTIDE, we performed computational analyses to study MACTIDE structure in solution.

**Figure 1.**
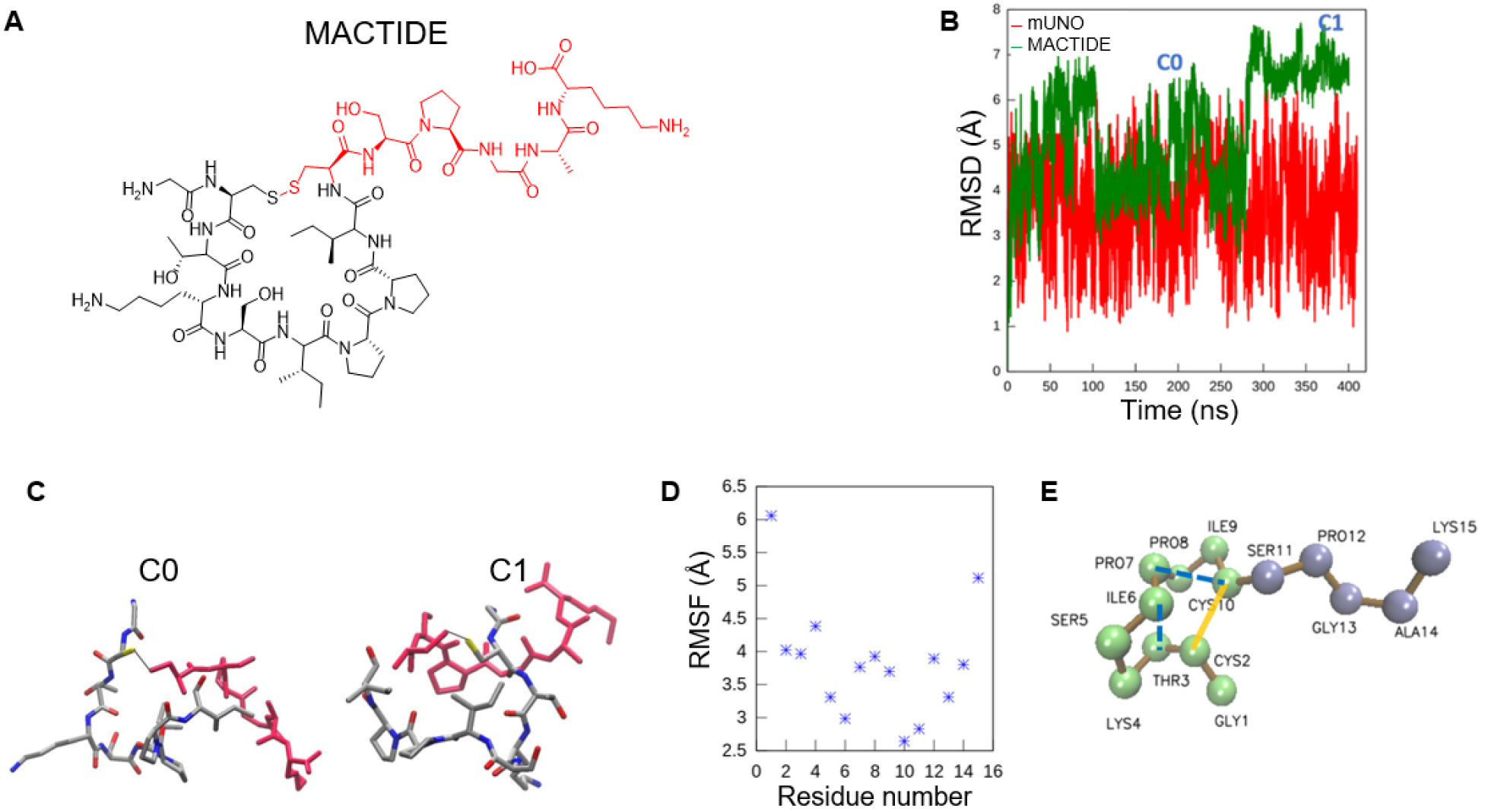
Design and molecular dynamics of MACTIDE. **(A)** MACTIDE structure with the CD206-binding motif mUNO in red. **(B)** RMSD to the average structure for heavy atoms of mUNO (red lines) and MACTIDE (green line), in two 400 ns molecular dynamics for mUNO and MACTIDE in solution. The two most populated conformations of MACTIDE, C0 and C1 are indicated in the figure. **(C)** structures of the C0 and C1 with the mUNO motif in red. **(D)** RMSF of each residue for MACTIDE in solution. **(E)** Intra hydrogen bonds in MACTIDE (dashed blue line), between Pro7-Cys10 with an occupancy of 75% and between Ile6-Thr3 with an occupancy of 10% (present in the C1 conformation).

An ensemble of MACTIDE conformations in solution was generated through an all-atom molecular dynamics simulation. The ensemble was clustered using a root mean square deviation (RMSD) criterion to group conformations with similar structural arrangements and identify the representative conformation. MACTIDE predominantly adopted two configurations, C0 and C1, with variations of about 1 Å from their centroid structure (Fig. 1B, green). In contrast, mUNO exhibited greater variation during simulations, with deviations reaching 3 Å from its average structure (Fig. 1B, red), indicating that MACTIDE is significantly more constrained than mUNO. The C0 and C1 configurations of MACTIDE were present in 44% of the entire trajectory and their structures are shown in Fig. 1C.

We also assessed the rigidity of MACTIDE by analyzing the root mean square fluctuation (RMSF) of each residue (Fig. 1D). Only the two terminal residues exhibit greater flexibility, while the rest of the peptide showed similar RMSF values. Additionally, we calculated the intra hydrogen bonds in MACTIDE (Fig. 1E, dashed blue lines). Both conformations exhibited a moderate hydrogen bond (mainly electrostatic) between Cys10 and Pro7, with a high occupancy of 75%. In the C1 conformation, there was an additional hydrogen bond between Ile6 and Thr3. The primary loop of the acyclic SFTI-1 only has the hydrogen bond Thr3-Ile9^12^, which was not observed in our structure, indicating that the structure of the primary loop in MACTIDE differs from that in acyclic SFTI-1.

### 2. Docking reveals high binding energy of MACTIDE to CD206 and ligand-induced conformational change

To estimate the binding site of MACTIDE, we used HPEPDOCK^15^, a hierarchical algorithm for blind and flexible peptide docking. The flexibility of CD206 was addressed through prior simulation of the receptor in solution, followed by clustering to identify the most populated receptor conformations (for more details see Materials and Methods). The best docking score was found for a receptor configuration where the alpha helix at CTLD2, defined by Thr360-Tyr373, is displaced downward by 3.8 Å, creating space for the peptide to accommodate (Fig. 2A). In the peptide-bound configuration, we noted slight differences in the CysR region and a closing of the V-shaped portion of the receptor compared to the crystal structure PDB: 5XTS (Fig. 2B). Interestingly, a similar ligand-induced conformational change in CD206 was observed when bound to mUNO^16^. The docking pose was located in the region between lectin domains CTLD1-2, the same region that binds mUNO^16^, with the peptide making closest contact with Asp273 and Thr324. Hydrogen bonds formed between G1-Thr 347 and P12-Gln 249 (Fig. 2C) and the mUNO motif (C10-K15) pointed toward the receptor. The HPEPDOCK docking score of MACTIDE was considerably higher than that of mUNO against the same receptor: -181.7 vs -115.0, respectively. Additionally, since G1 participated in a hydrogen bond with the receptor, we performed docking without it and found a lower docking score of -172.5, suggesting that G1 not only serves as a flexible spacer, but also contributes to binding. Based on this data, we decided to synthesize MACTIDE and evaluate it experimentally.

**Figure 2.**
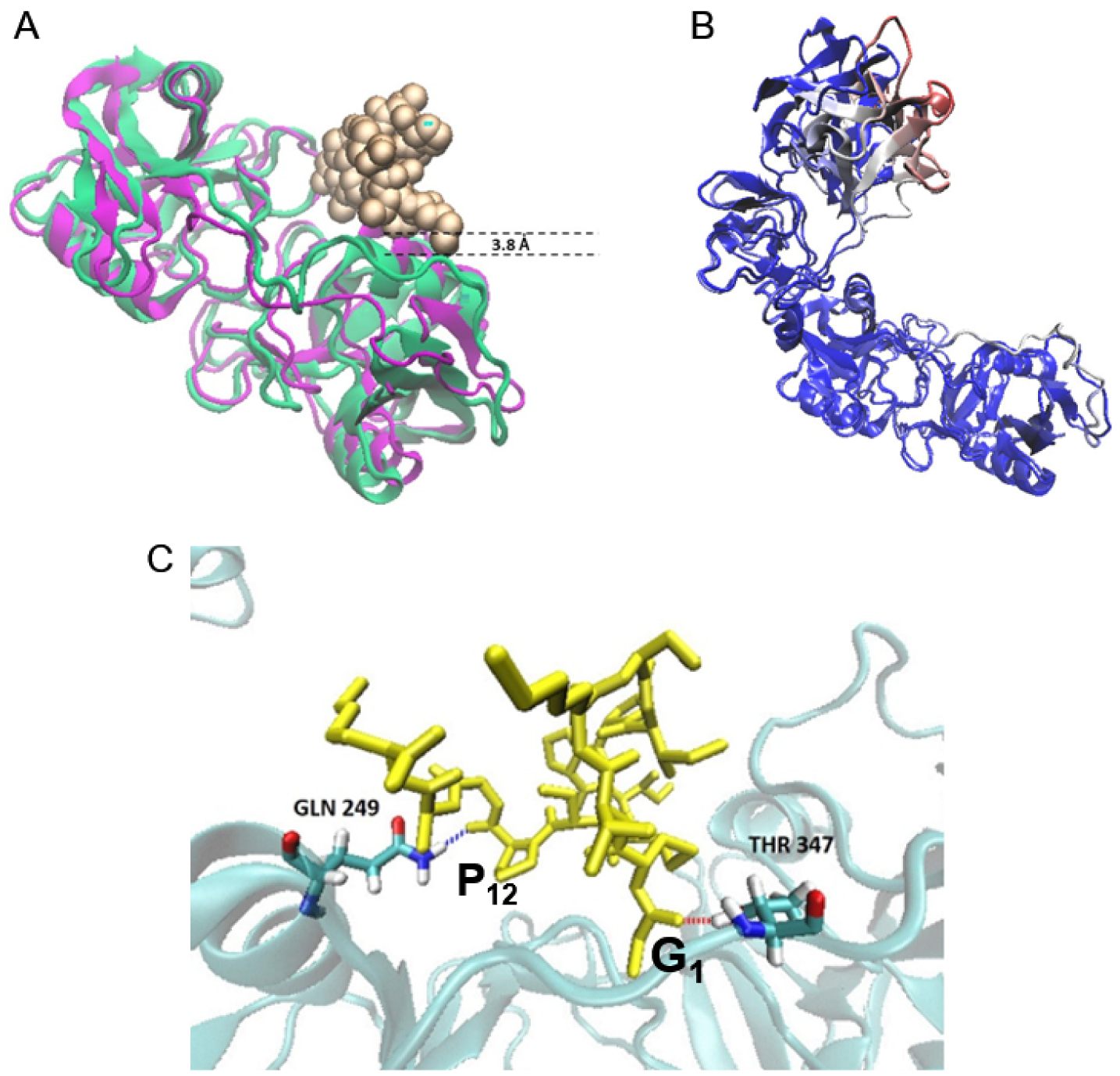
Docking shows high binding energy of MACTIDE to CD206 and ligand-induced conformational change. **(A)** CTLD2 domain for cluster node 2 (green) compared with cluster node 0 (purple). MACTIDE is represented as VDW golden spheres. A large alpha helix displacement of 3.8 Å was found in node 2 making space for the peptide to bind. **(B)** Node 2 compared with the crystal structure 5XTS and colored by RMSD. Significant differences (in red) were found at the CysR domain. **(C)** Hydrogen bonds formed at the docking pose.

### 3. MACTIDE shows enhanced affinity and proteolytic stability

To confirm the capacity of MACTIDE to bind to CD206, we evaluated binding to recombinant CD206 included in a viscoelastic film, using Quartz Crystal Microbalance (QCM). QCM is a powerful technique used to study label-free ligand-receptor interactions in solution^17,18,19,20^, wherein peptide binding and dissociation is sensed through a mass increase or decrease on a resonating crystal surface functionalized with the receptor. Here, we evaluated the mass changes of MACTIDE when binding and dissociating in solution to CD206 immobilized on a viscoelastic film^21^, deposited on a gold-modified quartz crystal resonating at 5 MHz. Recombinant CD206 was immobilized using layer-by-layer (LBL) self-assembly, an electrostatically-driven adsorption of charged polyelectrolytes on a layer of polyallylamine (PAH)^22^ ^23^ ^24^. These experiments showed that MACTIDE bound to PAH/CD206 (Fig. 3A, blue) and incompletely and slowly dissociated upon washing with PBS (Fig. 3A, blue arrow), whereas two control peptides did not bind to the same multilayer (Fig. 3A, gray and light green). Additionally, MACTIDE did not bind to the control multilayer PAH/BSA (Fig. 3A, dark green). The mUNO peptide also showed binding to the PAH/CD206 multilayer but with a complete and faster dissociation kinetics than MACTIDE upon washing with PBS (Fig. 3A, red), whereas the scrambled mUNO peptide did not bind to the same multilayer (Fig. 3A, cyan) and mUNO did not bind to the control multilayer PAH/BSA (Fig. 3A, brown). Fitting the association and dissociation curves revealed that the constant of association was slightly faster for mUNO (6.5 x 10^2^ M^-1^ s^-1^ versus 14 x 10^2^ M^-1^ s^-1^), but the main difference was in the dissociation constant, which was 30-fold slower for MACTIDE (2.5 x 10^-4^ s^-1^ versus 8.7 x 10^-3^ s^-1^), resulting in K_D_ = 0.38 µM for MACTIDE and K_D_ = 6 µM for mUNO.

**Figure 3.**
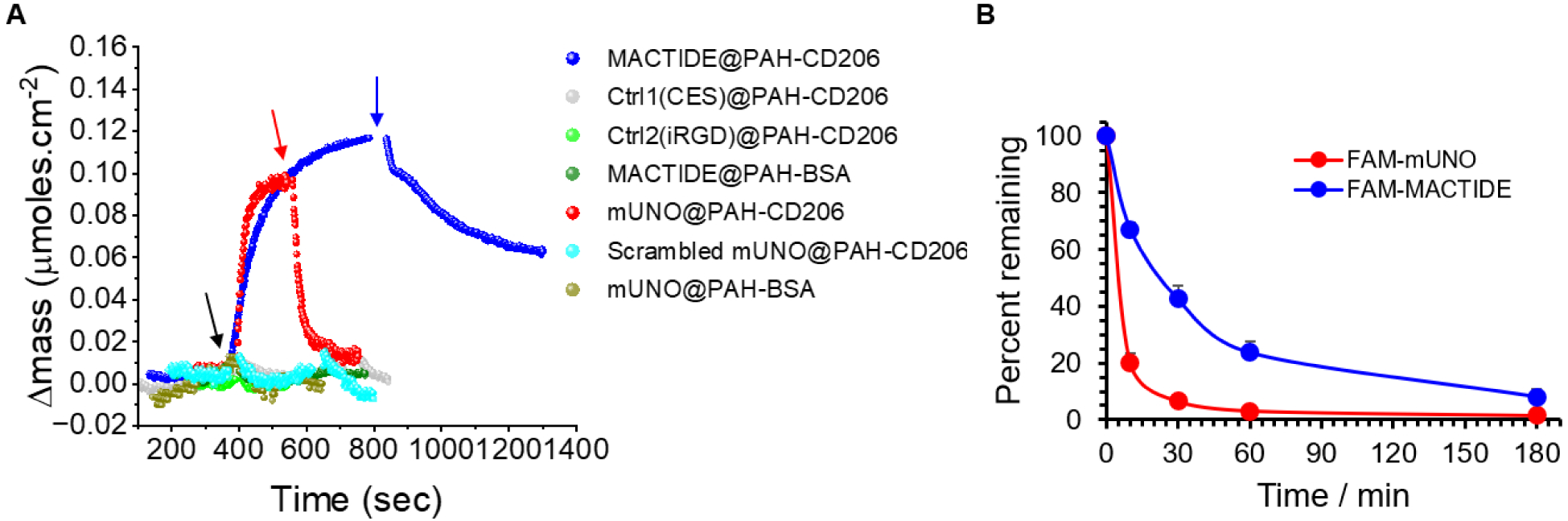
MACTIDE has higher affinity and proteolytic stability than mUNO. **(A)** QCM experiment of MACTIDE, mUNO and control peptides at final concentration of 10µM in PBS on multilayers of PAH/CD206 or control multilayers of PAH/BSA. The black arrow indicates when the peptides were added, blue and red arrows indicate when the washing step with PBS was started. **(B)** Integrity of FAM-MACTIDE and FAM-mUNO measured by LC-MS at different time points after incubation of peptides with lysate derived from a 4T1 tumor.

We next evaluated the stability of fluorescein (FAM)-MACTIDE against proteases from breast tumors. To this end we incubated both FAM-MACTIDE and FAM-mUNO with a tumor lysate obtained from 4T1 orthotopic TNBC tumors using different timepoints and evaluated the integrity of the peptide using liquid chromatography mass spectrometry (LC-MS). We observed that the half-life of FAM-MACTIDE in this tumor lysate increased 5-fold respect to FAM-mUNO (Fig. 3B).

### 4. FAM-MACTIDE targets CD206^+^ TAMs using different administration routes

To show that MACTIDE can be used to deliver a conjugated payload to CD206^+^ TAMs, we administered FAM-MACTIDE using different administration routes to mice bearing 4T1 tumors, which are highly infiltrated by CD206^+^ TAMs as we previously showed^2^. The targeting efficacy was evaluated by immunostaining of tumor sections for FAM and CD206.

FAM-MACTIDE showed CD206^+^ TAM targeting when administered intravenously (i.v.), whereas low CD206^+^ TAM targeting was observed for FAM-mUNO using the same route (Fig. 4A). With intraperitoneal (i.p.) administration, both FAM-MACTIDE and FAM-mUNO showed high CD206/FAM colocalization (80%) (Fig. 4B), but FAM-MACTIDE showed a 10-fold higher FAM intensity per CD206^+^ TAM. Importantly, FAM-MACTIDE also targeted CD206^+^ TAMs when administered orally (Fig. 4C). We also determined the blood half-life of FAM-MACTIDE using i.v. and i.p. administration (Fig. S1). As these values are the same as those obtained previously for FAM-mUNO^25^, we propose that the higher affinity and proteolytic stability of MACTIDE account for the homing differences observed between the two peptides. Importantly, we observed low liver accumulation with these administration routes for both peptides (Fig. S2).

**Figure 4.**
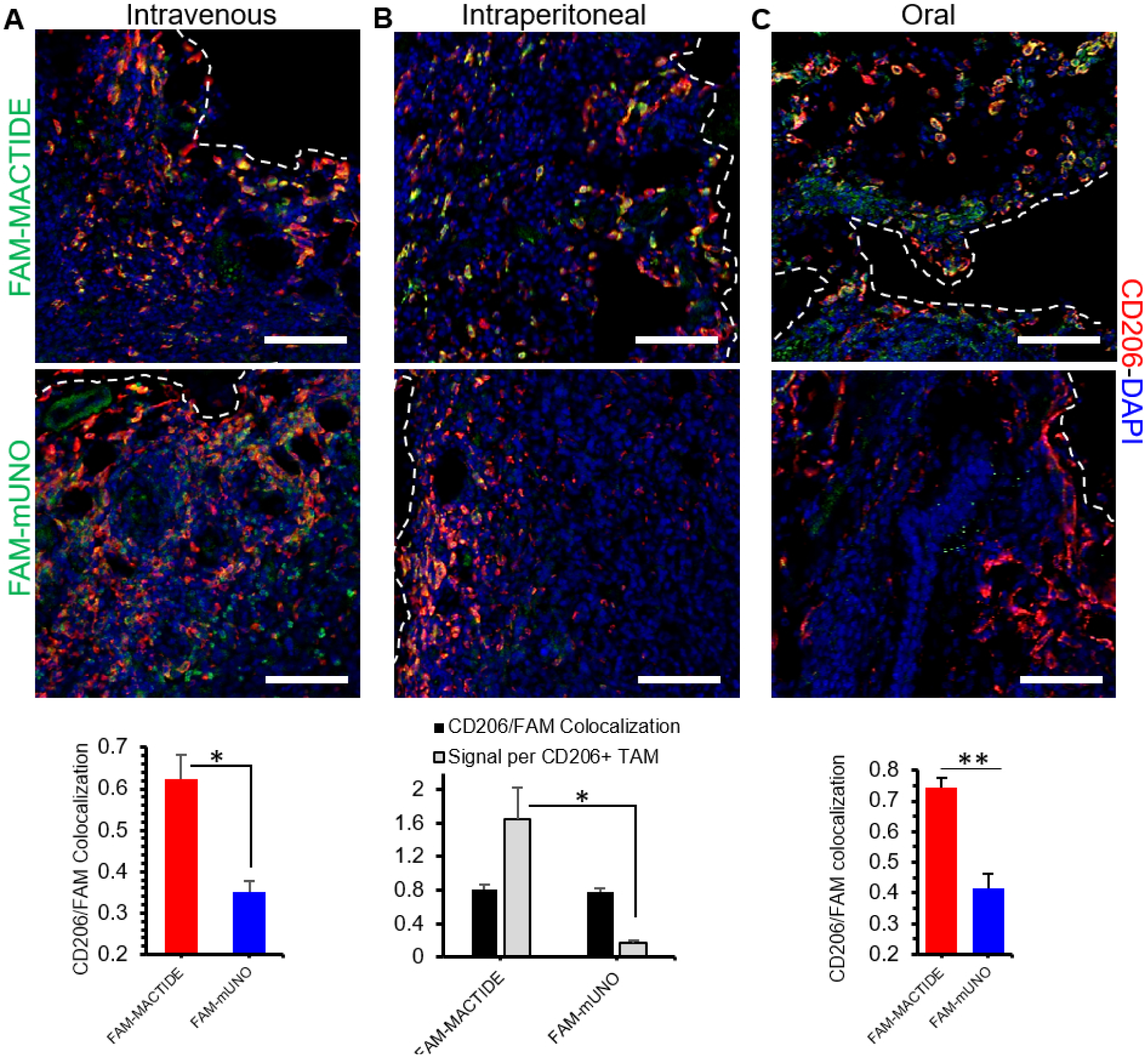
FAM-MACTIDE targets CD206^+^ TAMs using different administration routes. Thirty nmoles of FAM-MACTIDE or FAM-mUNO were administered i.p., i.v. or orally and left to circulate for 24 h. After 24 h, mice were sacrificed, and the organs were collected, fixed, cryoprotected, sectioned, and immunostained for FAM (shown in green) and CD206 (shown in red) **(A)**. The CD206/FAM colocalization indices were calculated from representative images from *n*=3 tumors, using Fiji (Mandler’s tM2 index) **(A, B, C)** and the signal intensity per CD206^+^ TAM was quantified using ImageJ for the i.p. administration **(B)**. Scale bars represent 100 μm. \**p* ≤ 0.05, \*\**p* ≤ 0.01 (Anova one way fisher LSD).

### 5. A MACTIDE-Verteporfin (MACTIDE-V) conjugate has light-dependent and light-independent *in vivo* activity in a breast cancer model

We next coupled MACTIDE to the photosensitizer Verteporfin, with the aim of depleting CD206^+^ TAMs using photodynamic therapy (PDT). To this end, carboxy-Verteporfin was coupled to the N-terminus of MACTIDE, a construct we refer to as “MACTIDE-V” (Fig. 5A). We performed *in vitro* photodynamic therapy with MACTIDE-V on primary human macrophages stimulated with IL-4 or LPS + IFNγ. Irradiated MACTIDE-V killed 80% of IL-4-stimulated macrophages and 40% of LPS + IFNγ-stimulated macrophages (we previously showed that these macrophages express CD206 at lower levels^26^). The non-irradiated MACTIDE-V showed no toxicity to these two cell types. Doxorubicin (DOX), a chemotherapeutic drug routinely used in clinic^27^, showed similar toxicity, but was independent of the irradiation, as expected. The control conjugates mUNO-V and CtrlPep-V showed no toxicity in any of the cells regardless of irradiation (Fig. 5B).

**Figure 5.**
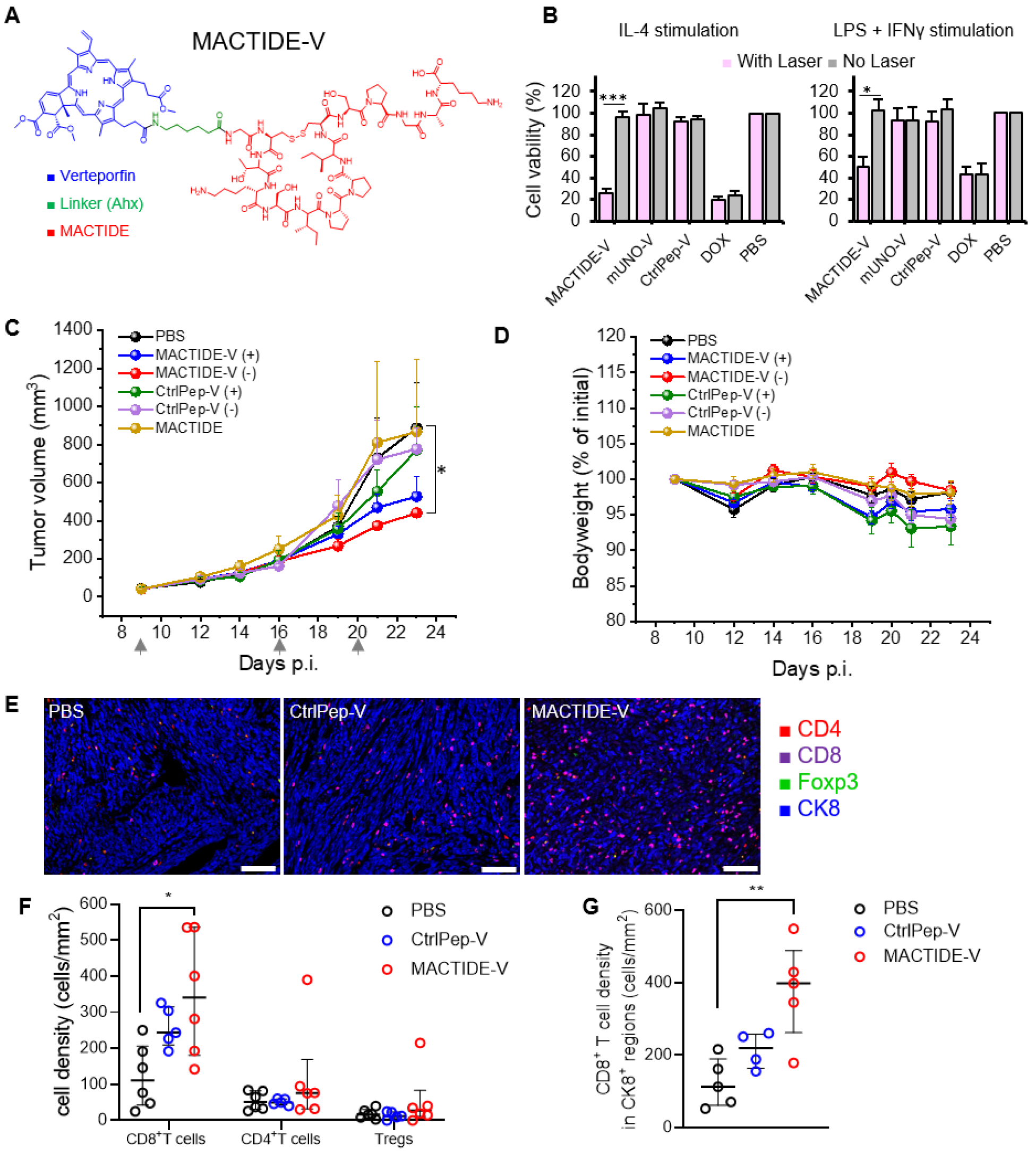
*In vitro* and *in vivo* photodynamic therapy with MACTIDE-V. **(A)** Structure of MACTIDE-V. **(B)** MACTIDE-V, CtrlPep-V and DOX were incubated with primary human macrophages (obtained from monocytes derived from human blood buffy coat) at 30 µM for one hour at 37°C, followed by 2 washes with medium. Then, the groups shown in black were irradiated with 10 J/cm^2^ (irradiance: of 170 mW/cm^2^, spot diameter: 0.5 cm). Then, the cells were incubated for 48 h and the cell viability was determined using the MTT((3-(4,5-Dimethylthiazol-2-yl)-2,5-Diphenyltetrazolium Bromide) assay. Here, the graph shows the average of *n*=6 experiments. (**C)** Treatment with MACTIDE-V and CtrlPep-V with and without laser (denoted by + and –, respectively), in mice bearing orthotopic 4T1 tumors (*n*=6/group) in Balb/C. Peptide-V conjugates were administered i.p. (indicated by gray arrows) at a dose of 30nmoles (1 mg/Kg in Verteporfin). For the irradiated groups, 4 h after administration of conjugates or PBS, mice were irradiated with 100 J/cm^2^, shown is primary tumor volume progression during treatment. Gray arrows indicate injection days. **(D)** Bodyweight during the treatment as % of the initial one. **(E)** Representative mIHC images of PBS, CtrlPep-V and MACTIDE-V-treated tumors, 20X magnification, scale bar=100 μm. **(F)** CD8^+^ T cells, CD4^+^ T cells and Treg density, *n*=6 mice/group, with every dot representing the mean cell density of each tumor obtained from 20 images. **(G)** CD8^+^ T cells density in CK8^+^-dense tumor regions. *n*=5 mice/group, with every dot representing the mean cell density of each tumor obtained from 20 images. Median ± interquartile range. One-way ANOVA with multiple comparisons, \**p* ≤ 0.05, \*\**p* ≤ 0.01.

Based on the consistent laser-induced and preferential toxicity of MACTIDE-V on IL-4-stimulated macrophages, we decided to evaluate its *in vivo* therapeutic efficacy on mice survival in the orthotopic 4T1 model. Although no significant differences were observed in mice survival (Fig. S3), MACTIDE-V followed by tumor irradiation produced a significant slowing down of tumor growth compared to the control groups PBS (+), CtrlPep-V (-), CtrlPep-V (+) and MACTIDE (-), where “+” denotes irradiated and “-” non-irradiated mice. Interestingly, non-irradiated MACTIDE-V-treated mice showed similar tumor volume reduction (Fig. 5C) and less effect on body weight (Fig. 5D) as irradiated MACTIDE-V.

Based on this, we decided to further investigate the inherent effect of MACTIDE-V without irradiation. To obtain further information on the effect of MACTIDE-V in this tumor model, we investigated T cell infiltration in treated tumors by multiplex immunohistochemistry (mIHC), comparing MACTIDE-V (-) treatment with PBS and CtrlPep-V (-) groups. mIHC analysis showed a significantly higher infiltration of CD8^+^ T cells in the MACTIDE-V group and no differences in the density of regulatory T cells (Tregs) (Fig. 5E, F). Importantly, the higher CD8^+^ T cell density was also observed in cytokeratin 8 (CK8^+^)-dense regions (Fig. 5G), highlighting the proximity of effector cells to cancer cells, crucial for anti-tumor cytotoxicity.

### 6. MACTIDE-V excludes YAP from the nucleus and increases phagocytosis and anti-tumoral gene expression in mouse bone marrow-derived macrophages

We speculated that the observed *in vivo* effect of MACTIDE-V could be mediated by the modulation of macrophage phenotype elicited by Verteporfin, a known inhibitor of the co-transcription factor Yes Associated Protein (YAP), which has been shown to prevent anti-inflammatory shift in macrophages^28^. We therefore analyzed the effect of MACTIDE-V on YAP localization and on phenotypic changes in mouse bone marrow-derived macrophages (BMDMs) *in vitro*. We first treated BMDMs with MACTIDE-V, MACTIDE or CtrlPep-V and evaluated changes in YAP localization 3 hours later using confocal microscopy. MACTIDE-V excluded YAP from the nucleus, whereas in the other experimental groups YAP was uniformly spread throughout the cell (Fig. 6A). These observations are consistent with the reported effect of Verteporfin of sequestering YAP in the cytoplasm^29^ ^30^.

**Figure 6.**
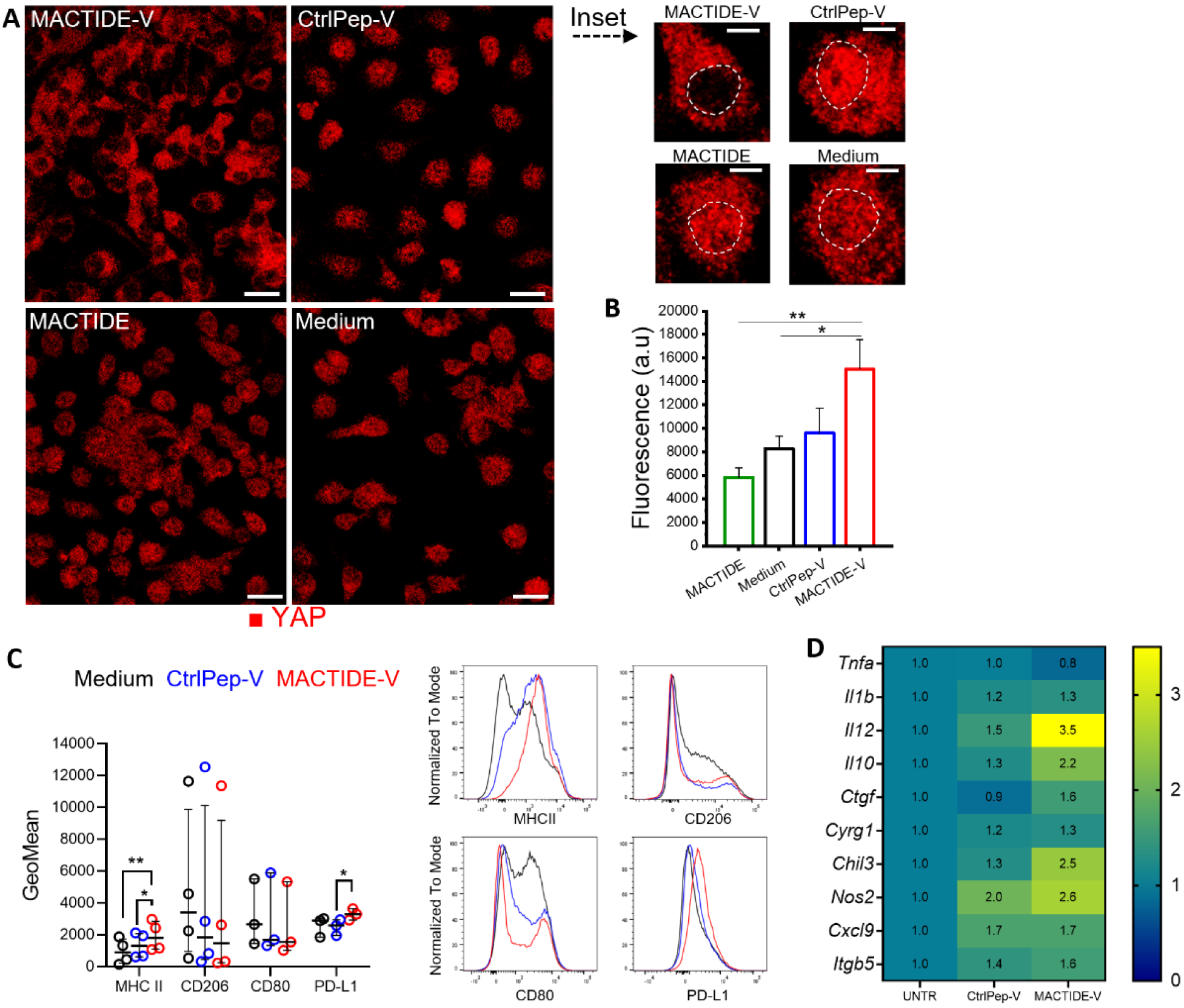
MACTIDE-V excludes nuclear YAP, increases phagocytosis, and MHCII expression in bone marrow derived macrophages. Day 5 BMDM were incubated 4 h with 10µM conjugates, washed, followed-up in medium. **(A)** YAP Immunofluorescence (shown in red) was analyzed 3 h after treatment and **(B)** phagocytosis of fluorescent *E.coli* particles at 48 h follow-up. **(C)** Flow cytometry on BMDMs 48 hours after treatment. *Left panel*, GeoMean of MHC II, CD206, CD80 and PD-L1 in Balb/c BMDMs. Median ± interquartile range. Repeated measures one-way ANOVA with multiple comparisons, for *n*=4 independent experiments. *p≤ 0.05, \*\**p* ≤ 0.01. *Right panel,* concatenated GeoMean histograms for MHC II in Balb/c BMDMs, *n*=4. **(D)** Heatmap of the mRNA expression of genes involved in the functional activation of BMDMs, YAP signaling and adhesion, measured by real-time PCR 48 hours after treatment in BALB/c BMDMs. *n*=3 independent experiments.

We then analyzed changes in BMDM phagocytosis 48 hours after treatments, as measured by the uptake of fluorescently labeled *E. coli* particles. MACTIDE-V significantly increased the phagocytosis relative to MACTIDE and untreated groups (Fig. 6B). As we observed no effects with free MACTIDE, we continued evaluating the effects of MACTIDE-V and CtrlPep-V only.

We subsequently analyzed the expression of different markers of BMDMs, both at the protein and the mRNA level, 48 hours after treatment with CtrlPep-V and MACTIDE-V. In Balb/C BMDMs, MACTIDE-V upregulated the expression of class II major histocompatibility complex (MHC II) and PD-L1 compared to CtrlPep-V and untreated conditions, with no significant changes in CD206 and CD80 expression (Fig. 6C and S6). Moreover, MACTIDE-V slightly affected the viability and differentiation of BMDMs, although this effect did not reach statistical significance (Fig. S4). At the mRNA levels, both conjugates induced an increase in *Il12*, *Il10, Chil3, Nos2* and *CXCL9* genes, more profound with MACTIDE-V treatment compared to CtrlPep-V (Fig. 6D). A similar tendency in the viability and phenotype after treatment was observed in BMDMs obtained from C57/Bl6 (B6) mice, where the upregulation of M1-related genes, such as *Il1b, Il12* and *Nos2,* was even more evident (Fig. S5).

### 7. MACTIDE-V has anti-tumoral and anti-metastatic effect in orthotopic 4T1.2

We then evaluated the therapeutic effect of MACTIDE-V on tumor progression and metastasis in the highly metastatic orthotopic 4T1.2 model^31^, better replicating TNBC in humans. When tumors reached 55 mm^3^ we began the treatment with MACTIDE-V or CtrlPep-V every other day until day 23, using the same dose of section 5. Four weeks after tumor injection, MACTIDE-V-treated mice showed significantly smaller tumor volumes compared to CtrlPep-V and PBS (Fig. 7A), as well as smaller endpoint tumor weights (Fig. 7B) and suppressed lung metastasis (Fig. 7C). No body weight loss was detected in any of the groups (Fig. S6).

**Figure 7.**
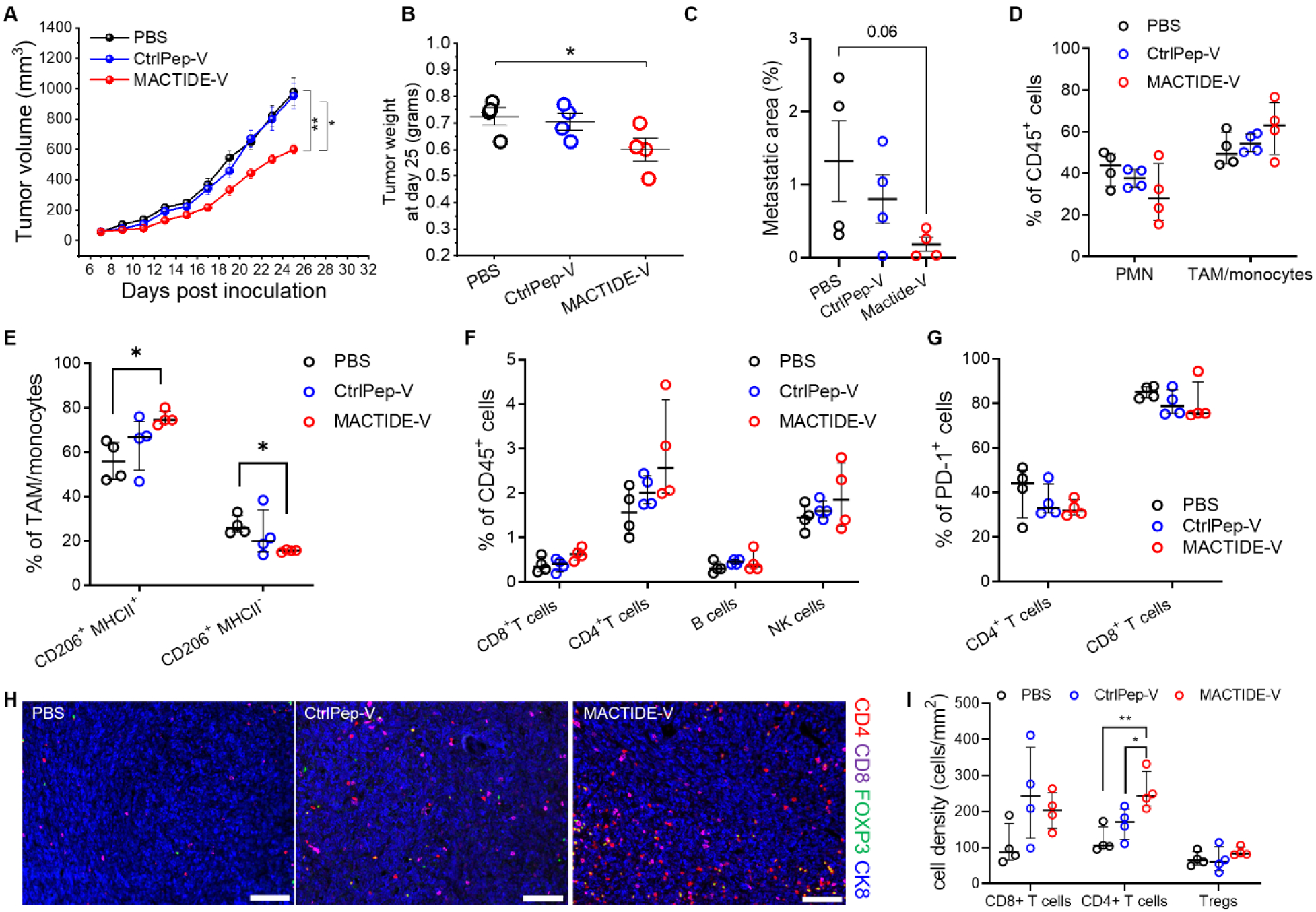
MACTIDE-V slows tumor growth and suppresses lung metastasis in orthotopic 4T1.2. 4T1.2 bearing mice were treated with 9 doses of CtrlPep-V, MACTIDE-V (1mg/Kg in Verteporfin per dose) or PBS i.p. every other day, while monitoring the primary tumor volume **(A)**. Mice were sacrificed on day 28, tumor weights were measured **(B)** and pulmonary metastasis areas were quantified from H&E sections **(C)**. **(D-G)** Tumor cell suspensions were analyzed by flow cytometry, *n*=4 per group. **(H)** Representative mIHC images of PBS, CtrlPep-V and MACTIDE-V treated tumors, scale bar=100 μm. **(I)** CD8^+^ T cells, CD4^+^ T cells and Treg density, *n*=6 mice/group, with every dot representing the mean cell density of each tumor obtained from 20 images. Median ± interquartile range. One-way ANOVA with multiple comparisons, * *p* ≤ 0.05, \*\**p* ≤ 0.01.

We then analyzed the immune cell populations of the tumor microenvironment by flow cytometry. MACTIDE-V did not reduce the percentage of CD206^+^ cells in population *in vivo* (Fig. S7A) and no effects were observed on the polymorphonuclear (PMN) or TAM/monocyte cell populations (Fig. 7D). MACTIDE-V increased the percentage of MHCII^+^CD206^+^ TAM/monocytes and decreased the MHC II^-^CD206^+^ subset (Fig. 7E).

In addition, MACTIDE-V treatment induced a rise in the proportion of tumor-infiltrating T cells and NK cells respect to CtrlPep-V and PBS, but no change in B cells (Fig. 7F). We also noticed a decrease in PD-1 expression in both CD8^+^ and CD4^+^ T cells in MACTIDE-V-treated tumors (Fig. 7G), also visible in CD25^+^ cells (Fig. S7B), possibly underlying a reduction in exhausted T cells^32^. Consistent with FC data, the mIHC analysis of 4T1.2 tumors revealed a significant increase in CD4^+^ T cell density in the MACTIDE-V-treated group compared to CtrlPep-V and PBS, a trend of increased CD8^+^ T cell density with both conjugates and no differences in Treg density (Fig. 7 H, I). These data show that MACTIDE-V is able to promote an anti-tumoral TAM/monocyte phenotype that is paralleled by an influx in the tumor mass of effector cells that might be less sensitive to PD-1 inhibition.

### 8. MACTIDE-V-treated tumors do not seem to benefit from concomitant anti-PD-1 blockade

To understand whether the treatment with MACTIDE-V could render 4T1.2 tumors more sensitive to immune checkpoint blockade, currently used in TNBC patients in combination with chemotherapy, we performed a treatment study with MACTIDE-V, anti-PD-1 or a combination of both, according to the scheme shown in Fig. 8A Both MACTIDE-V and anti-PD-1 blockade significantly reduced the primary tumor mass, but MACTIDE-V had a superior effect on reducing pulmonary metastases. Interestingly, no synergy was observed in the combination therapy in this setting (Fig. 8B-C). None of the experimental groups exhibited significant bodyweight loss (Fig. S8).

**Figure 8.**
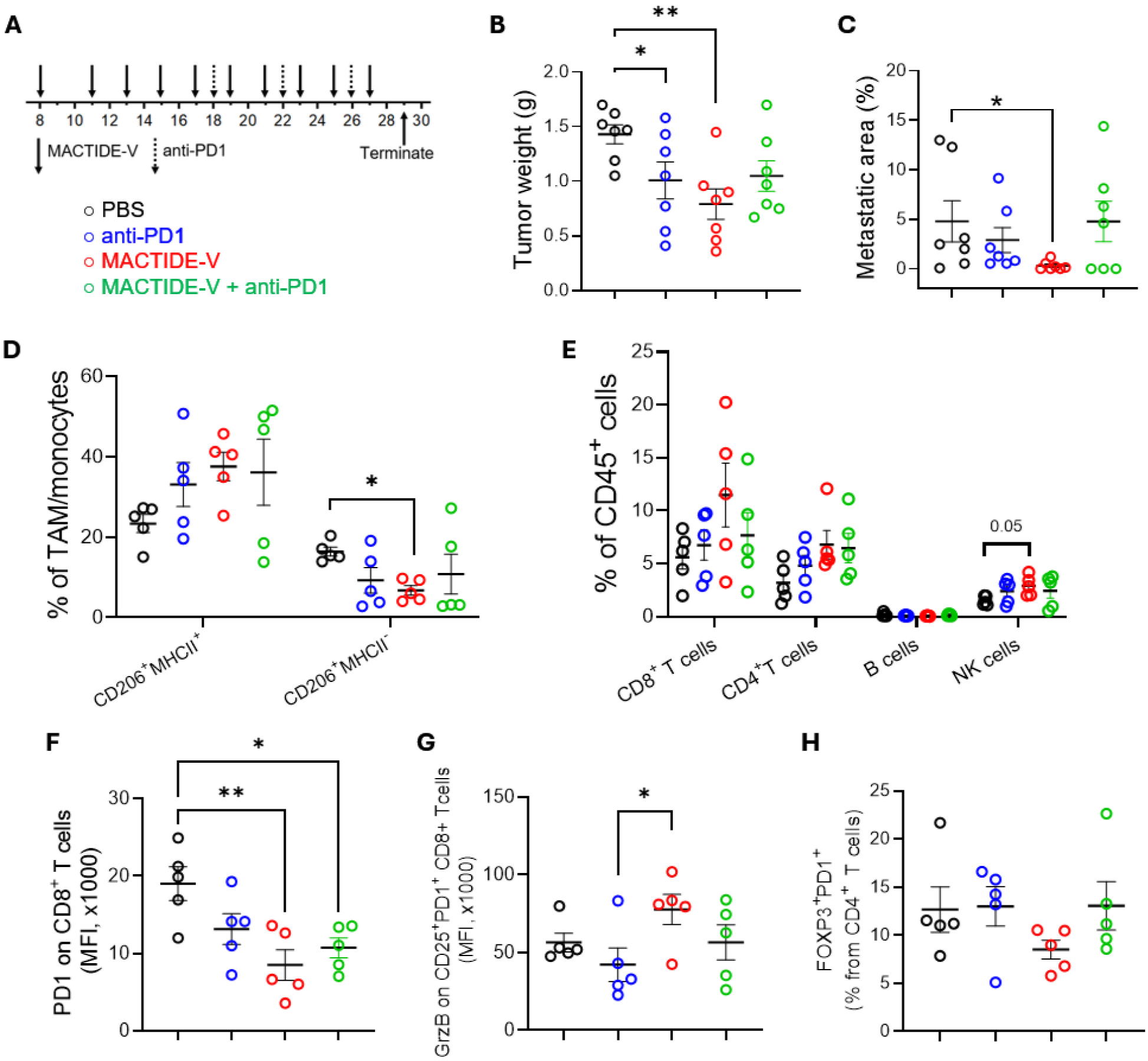
MACTIDE-V-treated tumors do not seem to benefit from concomitant anti-PD-1 blockade in orthotopic 4T1.2. 4T1.2 bearing mice were treated with 9 doses of CtrlPep-V, MACTIDE-V (1mg/Kg in Verteporfin per dose) or PBS i.p. every other day. Anti-PD-1 injections started 16 days post tumor induction, three injections of 200 µg each was given. Mice were sacrificed on day 29, their tumor weights analyzed **(B),** pulmonary metastases area quantified from H&E sections **(C)** and their tumors analyzed with flow cytometry **(D-I)**.

When we analyzed the composition of the tumor microenvironment by FC, we observed that MACTIDE-V elicited the highest increase in the CD206^+^MHCII^+^ population and highest decrease in the CD206^+^MHCII^-^ population of TAM/monocytes (Fig. 8D). MACTIDE-V also significantly increased the CD86^+^MHCII^+^ population and decreased the CD86^-^MHCII^-^ population of TAM/monocytes (Fig. S9).

MACTIDE-V elicited the highest increase in CD4^+^ T cells, CD8^+^ T cells and NK cells, although these differences reach statistical significance only for NK cells, with no differences in B cell infiltration (Fig. 8E). As already observed in the previous experiments, MACTIDE-V reduced the expression of PD-1 on CD8^+^ T cells (Fig. 8F) and also increased the expression of Granzyme B (GrzB) in CD25^+^PD-1^+^ CD8^+^ T cells (Fig. 8G). A trend in the decrease of the immunosuppressive FOXP3^+^PD-1^+^ population among CD4^+^ T cells was observed in the MACTIDE-V-treated group compared to the other experimental conditions (Fig. 8H).

These data suggest that MACTIDE-V can modulate TAM/monocytes by inducing antigen presentation and co-stimulation functions, resulting in an increase in effector cells with stronger cytotoxic potential in the tumor microenvironment.

## DISCUSSION

We here designed a CD206-targeting peptide, MACTIDE, of higher affinity, stability and oral activity over its predecessor peptide mUNO. The improved affinity of MACTIDE allowed us to develop a potent monovalent PDC, something which we did not achieve using mUNO as targeting peptide. Monovalent PDCs are attractive and translationally relevant drug candidates because they bypass the need for synthetically complex and multiparametric designs, such as nanoparticles or multivalent systems. Peptides and PDCs are becoming increasingly popular for cancer therapy, owing to their selectivity and high penetration in solid tumors^33, 34^. The other class of commonly used targeting ligands, antibodies, suffer from poor penetration in some solid tumors, and antibody-drug conjugates have shown toxicity-related limitations^35^.

Verteporfin (V), is a photosensitizer and inhibitor of the activation of YAP^36^. Our PDC, MACTIDE-V, was able to solubilize Verteporfin, which would otherwise need to be formulated in liposomes or be given with dimethyl sulfoxide as it is insoluble in water. Verteporfin, in its liposomal formulation Visudyne, is an approved drug for certain ophthalmic indications. In clinical trials for cancer therapy, Verteporfin is mostly used for photodynamic therapy (clinical trial identifiers: NCT03067051, NCT03033225, NCT04590664).

Here, our initial intention was to deplete TAMs using photodynamic therapy, but to our surprise, non-irradiated MACTIDE-V-treated mice experienced a similarly potent anti-tumoral effect *in vivo* than irradiated MACTIDE-V-treated littermates. Based on this observation, we further investigated the therapeutic effect of this PDC without irradiation, discovering that MACTIDE-V was altering the phenotype of TAMs, likely causing the anti-tumor effect.

Depletion of TAMs does not always result in strong anti-tumor effects in preclinical^37,38^ and clinical studies^39^ and might not be the most effective and safe strategy to modulate the tumor microenvironment as it could eliminate protective, sometimes anti-tumor, cell populations. On the other hand, the modulation of TAM phenotype appears to be a more promising approach, as suggested by other studies^40,41^ and supported by the results of this paper.

Our studies indicate that *in vivo* treatment with MACTIDE-V promoted a TAM/monocyte phenotype associated with phagocytosis and antigen uptake and presentation and also increased conventional CD4^+^ or CD8^+^ T cell infiltration without Treg increase, with induction of cytotoxicity markers in CTLs. MACTIDE-V also increased the number of NK cells, which is in line with studies showing that immunosuppressive TAMs also inhibit NK cells^42^. These immunostimulatory aspects of MACTIDE-V likely account for its anti-tumoral effect. The anti-metastatic effect of MACTIDE-V may be explained by the reduction of “M2-like” TAMs (here CD206^+^MHCII^-^), prominent culprits of metastasis in breast cancer^43^. A similar upregulation of pro-inflammatory and anti-tumoral markers was observed *in vitro* in BMDMs after MACTIDE-V treatment, further strengthening the idea that this PDC can modulate macrophages towards an M1-like or M1-M2 mix phenotype.

We observed that MACTIDE-V treatment did not reduce the CD206^+^ fraction of macrophages *in vitro* or *in vivo*. In addition to immunosuppressive M2-like TAMs, CD206 is also expressed by a subset of TAMs that participate in antigen uptake and presentation and stimulate anti-tumoral immunity^44^ and by phagocytic macrophages^45^. Interestingly, a correlation between the density of CD206^+^ TAMs and smaller tumor size and relapse-free survival has been reported in a cohort of TNBC patients^46^. For the above reasons, and supported by the findings of our paper, the approach of modulating the phenotype of CD206^+^ TAMs, instead of depleting these cells as we did in our previous study^26^, is likely a more effective therapeutic strategy for breast cancer.

Moreover, a combination treatment with MACTIDE-V and anti-PD-1 did not result in therapeutic synergy in our setting. Phagocytic macrophages were previously shown to interfere with T-cell based therapies. Arlauckas *et al*. found that phagocytic TAMs took up the administered anti-PD-1 from the surface of the T-cells^47^ and Yamada-Hunter *et al*. showed that activating macrophage phagocytosis can lead to TAMs phagocytosis of chimeric antigen receptor (CAR)-T cells^48^. In our study we noticed that PD-1 was downregulated after MACTIDE-V treatment, making us hypothesize that the concomitant administration of anti-PD-1 might dampen the effects of this immunotherapeutic drug. These results highlight the challenges in designing “TAM-aimed” together with “T cell-aimed” therapies and the need for further studies to fully comprehend these complexities and optimize treatment schedules.

In practical terms however, our results support safety and a consistent anti-tumor effect of MACTIDE-V monotherapy. Given the strong anti-metastatic effect of MACTIDE-V, one would envision using MACTIDE-V as a neo-adjuvant agent in metastatic breast cancer prior to resection of the primary tumor, or as adjuvant agent together with chemotherapy, prior or subsequent (but not concomitant) to immunotherapy.

MACTIDE-V represents a valuable peptide-drug conjugate for reprogramming TAM/monocytes and the MACTIDE peptide represents a potent tool to target CD206^+^ TAMs, even using oral administration. Further studies will need to assess MACTIDE-V in other tumor models, further dig into its mechanism of action and explore the oral route of administration.

## MATERIALS AND METHODS

### Peptides and peptide conjugates

Peptides were synthesized on solid phase. FAM and Vert denote carboxyfluorescein and Verteporfin respectively and they were coupled to the N-terminus of peptides via their carboxylic acid, spaced via an aminohexaonic acid linker (Ahx). Peptides and FAM-peptide conjugates were purchased from Lifetein LLC, TAG Copenhagen or synthesized at the peptide synthesis core facility of CNB-CSIC, Madrid, Spain.

Verteporfin-peptide conjugates were prepared in the Peptide Synthesis Unit (U3) at IQAC-CSIC (https://www.nanbiosis.es/portfolio/u3-synthesis-of-peptides-unit/). The peptide moiety was synthesized on a microwave-assisted peptide synthesizer (Liberty Blue,CEM), using Rink amide Protide resin (0.56 mmol/g, CEM) as a solid support and a Fmoc/tBu strategy. Diisopropylcarbodiimide (DIC) and Oxyme were used as coupling reagents. After completion of the peptide moiety, verteporfin (2 eq.) was manually introduced. Vert-peptide conjugates were released from the solid support by treatment with TFA: CH_2_Cl_2_:TIS (95:2.5:2.5, v/v/v) for 1.5 h. The solvent was then evaporated under vacuum, and the peptide conjugates were precipitated with cold diethyl ether and centrifugated. Then, the liquid was decanted and the solid dissolved in a mixture of H_2_O:CH_3_CN (1:1, v/v) and lyophilized. To generate the MACTIDE-V with the disulphide bridge, a 1mM solution of the linear precursor of MACTIDE-V in H_2_O:CH_3_CN (1:1, v/v) was prepared and the pH was adjusted to 8 with a solution of 20% NH_4_Cl in H_2_O. The evolution of disulphide formation was monitored by HPLC and was completed after 12 h. MACTIDE-V was purified by semipreparative HPLC with a XBridge Peptide BEH C18 OBD Prep column (130 Å, 5 μm, 19 x100 mm), using H_2_O (1% CF_3_COOH) and CH_3_CN (1% CF_3_COOH) as eluents. Final pure peptide conjugates were analyzed and characterized by HPLC and HPLC-MS.

Peptides and conjugates used were the following:

| Notation used | Peptide | Notes |
| --- | --- | --- |
| FAM-MACTIDE | FAM-Ahx-G(CTKSIPPIC)SPGAK-OH | Disulfide: C2-C10 |
| MACTIDE | G(CTKSIPPIC)SPGAK-OH | Disulfide: C2-C10 |
| mUNO | CSPGAK-OH |  |
| Ctrl1 | Ac-(CESPLLSEC)-NH<sub>2</sub> | Disulfide: C1-C9 |
| Ctrl2 | Ac-(CRGDKGPDC)-NH<sub>2</sub> | Disulfide: C1-C9 |
| CtrlPep-V | Vert-Ahx-AKPCGS-OH |  |
| MACTIDE-V | Vert-Ahx-G(CTKSIPPIC)SPGAK-OH | Disulfide: C2-C10 |

### Molecular dynamics

MACTIDE was constructed with tLeap from Amber Package^49^, modelling interactions using the ff14SB amber forcefield^50^ It was solvated with TIP3P water^51^, Cl-ions were added to neutralize the net charge. Three independent molecular dynamics simulations were performed, each 400 ns long. Simulations were conducted using the Amber18 Package with the following protocol: first a minimization was performed to relax clashes with the steepest descent method combined with the conjugate gradient. Next, temperature and pressure were included with short simulations using NVT and NPT ensembles and once the systems were equilibrated the production runs were started. The time step used was 2 fs. Electrostatic interactions were treated using particle-mesh Ewald (PME)^52^ with a cut-off of 10 Å. Temperature was regulated using Langevin dynamics^53^ with a collision frequency of 2 ps^-1^. Simulation trajectories were saved every 10 ps. Representative conformations were extracted from the trajectories by performing a clustering analysis, using a hierarchical agglomerative approach. MACTIDE was solvated with water and two independent molecular dynamics simulations were performed, each 400 ns long, to construct an ensemble of configurations.

### Peptide docking analysis

Docking was conducted using HPEPDOCK^54^, which involves global sampling of binding orientations along the receptor surface. This algorithm accounts for peptide flexibility by generating an ensemble of conformations, which are then globally docked against the entire protein. To address receptor flexibility, we used an ensemble of receptor conformations from a previously generated trajectory (3). These conformations were clustered based on an RMSD criteria. The five most populated receptor configurations were selected and used as coordinates for the receptor in each docking calculation.

### Peptide stability in tumor lysate

For peptide stability measurements, 200 µl of freshly prepared tumor lysate was mixed with 50 µl of PBS (as control), 30 µM of FAM-mUNO, or 30 µM of FAM-MACTIDE and incubated at 37 °C. Forty µl aliquots were taken at the 0, 10, 30, 60, 180 and 1440 minutes. Eighty µl of methanol was added to each aliquot and it was immediately stored at -80 °C until analysis later on the same or following day. For analysis, the samples were centrifuged at 21 000 × g for 10 min at +4 °C. The supernatant was transferred into liquid chromatography (Agilent 1200 series) autosampler and maintained at +4 °C. Ten µl was injected and separation was achieved with a C18 column (Kinetex 2.6 µm EVO C18 100×4.6 mm, Phenomenex). The chromatography gradient started with 5 min 5 % acetonitrile in water, followed by linear increase to 100 % acetonitrile in 20 min and finally 20 min isocratic flow of 100 % acetonitrile. Both eluents contained 0.2 % formic acid. Enhanced resolution scan (Qtrap 4500, Sciex) for peptides with 1, 2 or 3 charges was done. Additionally, all m/z values in the 50-2000 range were scanned for degradation product search. Potential product signals were subjected to fragmentation analysis. Statistics were done using GraphPad Prism 5.0.

### Peptide binding studies using Quartz Crystal Microbalance

The Quartz Crystal Microbalance (QCM) used was QCM200 system from Stanford Research Systems (Sunnyvale, CA, USA). The quartz crystals used have 5MHz resonant frequency and are deposited with a layer of Cr/Au (Cat # O100RX1, p/n 6-613, Stanford Research Systems). The crystal was mounted on the cell, washed with isopropanol, ethanol and mQ water. Then, it was incubated with 20 mM Mercapto-propanesulfonate (MPS, Cat #251682, Sigma-Aldrich) in 10 mM H_2_SO_4_ for 30 minutes, washed with mQ and then incubated with a 10 µM (in monomer) of Polyallylamine hydrochloride, Mw: 50000, PAH, Cat # 283223, Sigma-Aldrich) in mQ at pH 8.5 during 10 min and later washed with mQ. Then, a baseline with 500 µL of mQ at pH 8 was recorded, the measurement was paused and then 5 µL of a 1 mg/mL solution in PBS was added (human recombinant CD206, Cat # 2534-MR-050/CF from R&D systems, reconstituted with 50 µL of mQ); final concentration of CD206 in the cell: 0.01 mg/mL. Of note, the isoelectric point of CD206 is 6.3. For PAH-BSA multilayer, Bovine Serum Albumin (BSA, Cat # A7906, Sigma-Aldrich) was deposited, after measuring a baseline in mQ, at a concentration of 0.01 mg/mL in mQ for 10 minutes, and then washed with mQ. For peptides, 500 µL of PBS was placed in the cell and the baseline was recorded, the measurement was then paused and 50 µL of a 100 µM solution of peptides in PBS was gently deposited on the cell and the measurement was resumed. Then, the measurement was paused, the solution removed and replaced with 500 µL of new PBS and the measurement resumed. The association and dissociation curves were fitted using TraceDrawer software (Ridgeview Instruments AB), to obtain the association constant k_a_, the dissociation constant k_d_ and the affinity constant K_D_.

### Cell culture and experimental animals

4T1 and 4T1.2 cells were both purchased from ATCC. 4T1 cells were grown in RPMI1640 medium (Gibco™, catalog no. 72400-021) supplemented with 10% v/v fetal bovine serum (FBS, Capricorn Scientific, catalog no. FBS-11A) and 100 IU/mL penicillin/streptomycin (Pen/Strep, Capricorn Scientific, catalog no. PS-B) at +37 °C in the presence of 5% CO_2_. 4T1.2 cells were grown in AlphaMEM (Gibco™, catalog no. 12571063) supplemented with 10% v/v FBS and 100 IU/mL Pen/Strep at +37 °C in the presence of 5% CO_2_.

All animal experiments were performed on 8-12-week-old female Balb/c mice and were approved by the Estonian Ministry of Agriculture (project no. 197). All methods were performed in accordance with existing guidelines and regulations.

### *In vivo* biodistribution studies

Orthotopic tumors were induced by injecting subcutaneously 1 x 10^6^ 4T1 cells in 50 µL of PBS (Lonza, catalog no. 17-512F) into 4^th^ mammary fat pad of 8-12-week-old female Balb/c mice. When tumors reached approximately 100 mm^3^, thirty nmoles of FAM-MACTIDE or FAM-mUNO was injected i.p., i.v. or through oral gavage and circulated for 24 h. Then, mice were sacrificed by anesthetic overdose and cervical dislocation, organs and tumors were collected and fixed in cold 4% paraformaldehyde (PFA) in PBS at +4 °C overnight followed by washing in PBS at RT for 1 h after which 15% w/v sucrose was added for 24 h. Finally, 30% sucrose was added overnight, cryoprotected tissues were frozen in Optimal Cutting Temperature (OCT; Leica, catalog no. 14020108926) medium and stored at -80 °C for long term or at -20 °C for short term. Blocks were cryosectioned at 10 µm thickness on Superfrost+ slides (ThermoFisher Scientific, catalog no. J1800AMNZ) and stored at -20 °C or used instantly. Immunofluorescence staining was performed as described previously^5^. CD206 was detected using rat anti-mouse CD206 (dilution 1/200, Bio-Rad, catalog no. MCA2235GA) and Alexa Fluor 647 goat anti-rat antibody (dilution 1/300). FAM was detected using rabbit anti-mouse FAM (dilution 1/100) and Alexa Fluor 546 goat anti-rabbit antibody (dilution 1/200). Slides were counterstained using 4,6-diamidino-2-phenylindole (DAPI, 5 μg/mL in PBS, Sigma-Aldrich, catalog no. D9542-5MG). Slides were mounted using mounting medium (Fluoromount-G Electron Microscopy Sciences, catalog no. 17984-25) and imaged using Zeiss confocal microscope (Zeiss LSM-710) and 20 x objective. The colocalization analysis was performed using “Coloc2” plugin in the Fiji program using Mandler’s tM2 index. Values were obtained from at least three individual images per mouse per group and their average values were plotted. The FAM mean signal per CD206^+^ cell analysis was measured using Fiji, taking the mean FAM signal, and dividing it by the number of CD206^+^ cells. Average values were obtained from four images per mouse for *n*=3 mice.

### *In vitro* photodynamic therapy

Human peripheral mononuclear cells (PBMCs) were purified from human blood buffy coat following the protocol described previously^26^. Briefly, we used Ficoll Paque Plus (GE Healthcare, catalog no. 17-1440-02) reagent and CD14^+^ microbeads (MACS Miltenyi Biotec, catalog no. 130-050-201), seeded 1.2 x 10^5^ cells in 50 µL of RPMI1640 medium on FBS-coated 96-well plate. For optimal cell attachment and polarization, macrophage colony stimulating factor (M-CSF, 50 ng/mL, BioLegend, catalog no. 574802) was added. Then, to obtain M2 resembling phenotype, monocytes were stimulated with IL-4 (50 ng/mL, BioLegend, catalog no. 574002). 50 µL of medium containing M-CSF and IL4 was replenished every other day for 7 days. To obtain M1 resembling phenotype, monocytes were stimulated with M-CSF for 6 days (50 ng/mL), 50 µL replenished every other day after which lipopolysaccharide (LPS, 100 ng/mL, Sigma-Aldrich, catalog no. L4391) and IFNγ (20 ng/mL, BioLegend, catalog no. 570202) were added and incubated overnight. All incubations were done at +37 °C. On day 7, 30 µM of MACTIDE-V, mUNO-V, CtrlPep-V, DOX or PBS was added, and cells were incubated for 60 min at + 37 °C after which cells were washed with medium and 100 µL of new RPMI without phenol red (Gibco™, catalog no. 11835030) was added. *n*=3 wells/group from *n*=6 donors. Cells were then irradiated using a NIR laser source for PDT from Modulight Inc (ML6500, 2W, 689nm) and an optical fiber with frontal diffuser (SMA905, Modulight), dose 10 J/cm^2^ and spot size 0.5 cm. As Verteporfin is light sensitive, everything was performed in dark. To keep conditions the same, the plate that did not receive irradiation was also kept open for the same about of time. After irradiation, cells were incubated at +37 °C for 48 h. To analyze cell death, 10 µL of 4,5-Dimethylthiazol-2-yl)-2,5-Diphenyltetrazolium Bromide (MTT, 5 mg/mL) in PBS was added to the cells and incubated at +37 °C up to 90 min. Crystal formation was monitored every 20-30 min to not oversaturate the OD values. Then, medium was removed carefully and 100 µL of isopropanol was added to each well and plate was shaken until all crystals were dissolved. Absorbance was read at 570 nm using a plate reader (Tecan Sunrise) and corresponding program (Magellan 7).

### *In vivo* photodynamic therapy in orthotopic 4T1

Orthotopic tumors were induced by injecting 5x10^4^ 4T1 cells in 50 µL of PBS subcutaneously into 4^th^ mammary fat pad of 8-12-week-old female Balb/c mice. When tumors reached approximately 40 mm^3^, mice were sorted into groups: MACTIDE-V (+), MACTIDE-V (-), CtrlPep-V (+), CtrlPep-V (-), MACTIDE (-) and PBS (+). Tumors were measured using a digital caliper (Mitutoyo) and volume calculated using (W^2^ x L)/2 formula, where W is the width of a tumor and L is the length.

Each group had six mice. First intraperitoneal injection (30 nmoles) was carried out on day 9 post tumor induction. Four hours post injection, tumor area was shaved to lessen laser scattering and tumors were irradiated with 100 J/cm^2^ using the NIR laser source for PDT described above. Each tumor was measured, and spot size adjusted every time, keeping the radiation dose always constant. Irradiation of the tumors was done in complete darkness under anesthesia. Mice recovered on a heating pad and eye drops were applied to avoid eye dryness. All mice, whether irradiated or not, were anesthetized to keep the handling conditions the same. Mouse bodyweights and tumor volumes were monitored every other day. The sacrifice of mice began on day 23 based on their tumor sizes (over 1500 mm^3^) or cachexia. The final mice were sacrificed on day 36 post tumor induction. Tumor volume curves are shown until day 23, when mice elimination began, and the sample number became too small for statistical comparison. Survival was analyzed using GraphPad Prism to plot Kaplan-Meier survival curves and perform Mantel-Cox test for statistical analysis. Tumors taken on similar days were analyzed using mIHC.

### Multiplex immunohistochemistry (mIHC)

The staining of 4 µm-thick formalin-fixed paraffin-embedded (FFPE) tissue sections for mIHC analysis was carried out using the tyramide signal amplification-based Opal method (Akoya Biosciences) on a Leica BOND RX automated immunostainer (Leica Microsystems). For each staining cycle, FFPE slides were deparaffinized and subjected to heat-induced epitope retrieval at 97 ° or 100 °C using BOND epitope retrieval solutions ER1 or ER2 (Leica Biosystems). The tissue sections were incubated with a blocking solution for 10 minutes, then incubated for 30 minutes with primary antibodies listed in the table below.

| **Antigen** | **Clone** | **Reference** | **Dilution** | **Incubation time** | **Secondary antibody** | **OPAL** |
| --- | --- | --- | --- | --- | --- | --- |
| CD4 | 4SM95 | Invitrogen 14-9766-82 | 1:150 | 30' | anti-rat | 480 |
| CD8 | 4SM15 | Invitrogen 14-0808-82 | 1:150 | 30' | anti-rat | 620 |
| Ly6G | EPR 22909-135 | Abcam ab238132 | 1:500 | 30' | anti-rabbit | 570 |
| Fibronectin | polyclonal | Abcam ab2413 | 1:250 | 30' | anti-rabbit | 690 |
| FOXP3 | 221D | Abcam ab253297 | 1:200 | 30' | anti-mouse | 650 |
| Cytokeratin 8 | EP1628Y | Abcam ab53280 | 1:200 | 30' | anti-rabbit | 780 |

The slides were then incubated for 10 minutes with the horseradish peroxidase-conjugated secondary antibodies Opal polymer HRP mouse-rabbit (Akoya Biosciences), ImmPRESS® HRP Goat Anti-Rat IgG HRP-polymer or ImmPRESS® HRP Goat Anti-Rabbit IgG HRP-polymer (Vector Laboratories). After washing, the slides were incubated for 10 minutes with the tyramide signal amplification-conjugated fluorophores Opal-480, 570, 620, 650, 690 and 780 (Akoya Biosciences) and with the spectral DAPI (Akoya Biosciences) as a nuclear counterstain. After washing the slides were mounted using the ProLong diamond antifade mountant (Invitrogen, ThermoFisher Scientific) and imaged using a Mantra2 quantitative pathology workstation (Akoya Biosciences) at X20 magnification. At least 20 fields were acquired for each slide. The spectral unmixing of the images was performed with InForm 2.6 Image Analysis Software (Akoya Biosciences) and the analysis of cell density with QuPath v0.5.1. Fibronectin and Ly6G staining (not shown) were used to better identify non-necrotic tumor regions.

### RNA extraction, retro transcription and real-time PCR

For RNA extraction, BMDMs were detached, pelleted and resuspended in 1 mL of TRIzol® reagent (Cat# 15596018, Invitrogen, ThermoFisher Scientific) and stored at -80°C until the RNA extraction. Total RNA was extracted following TRIzol® manufacturer’s protocol and RNA was quantified using NanoDrop One (Thermofisher Scientific). 1000 ng of RNA were used for cDNA retrotranscription using the High-capacity RNA-to-cDNA™ Kit (Cat# 4387406, Applied Biosystems, Thermofisher Scientific) following manufacturer’s instructions. 20 ng of cDNA per well were amplified in 20 μL using the TaqMan® Universal PCR Master Mix (Cat# 4364340) and TaqMan® assay primers and probes (Cat# 4448892 or 4453320, Applied Biosystems, Thermofisher Scientific). The reaction was performed in MicroAmp® Optical 96-well reaction plate (Cat# N8010560, Applied Biosystems, Thermofisher Scientific) and analyzed on a QuantStudio™ 7 Pro Real-Time PCR System. All the samples were amplified in duplicates and data were analysed using the ΔΔCt method using *Gapdh* as housekeeping gene. The following Taqman® Gene Expression Assays were used:

| Gene | Taqman assay ID |
| --- | --- |
| *Tnfa* | Mm00443258_m1 |
| *Il1b* | Mm00434228_m1 |
| *Il12* | Mm00434174_m1 |
| *Il10* | Mm00439614_m1 |
| *Ctgf* | Mm01192933_g1 |
| *Cyrg1* | Mm00487498_m1 |
| *Chil3* | Mm00657889_mH |
| *Nos2* | Mm00440485_m1 |
| *Cxcl9* | Mm00434946_m1 |
| *Itgb5* | Mm00439825_m1 |
| *Gapdh* | Mm99999915_g1 |

### Mouse BMDM differentiation and treatment with conjugates

Bone marrow cells were isolated from 8-12 weeks old female Balb/c mice by flushing the femur and tibia bones with medium using a 25G needle. Red blood cells were lysed (RBC lysis buffer for mouse, ThermoScientific, cat # J62150.AK) and cell suspensions were washed, resuspended in complete medium and filtered with a 100 μm cell strainer. Macrophages were obtained by culturing the bone marrow cells for 5 days at a density of 2x10^5^ cells/cm^2^ in RPMI1640 (Gibco, cat # 11875-093) supplemented with 10% (v/v) FBS and 1% penicillin/streptomycin, and 100 ng/ml M-CSF (BioLegend, cat # 576406) at 37°C, 5% CO_2_. Culture media was refreshed every 2-3 days by substituting half of the medium with fresh one containing M-CSF. On day 5, cell culture medium was removed and replenished with an equal volume of medium containing the peptide conjugates at the final concentration of 10 μM. After an incubation of 4 hours at 37°C, the peptide conjugates were removed by a washout and BMDMs were incubated for 48 hours at 37°C with fresh medium.

### YAP immunofluorescence

For YAP immunostaining, cells were cultured on Ibidi plate (Ibidi, cat# 80821) previously coated with FBS and treated with conjugates as previously described. 3 h after peptide conjugates were removed, the medium was also removed, and cells were washed with PBS and fixed with PFA (ThermoScientific, Cat # J61899.AP). Fixed cells were then washed with PBS, permeabilized with PBS-0.2% Triton-X100 (v/v) and washed with PBS 0.05% Tween-20 (PBS-T) (v/v), before being incubated with blocking buffer (5% BSA (w/v), 5% FBS (v/v) in PBS-T). After blocking, cells were incubated overnight at +4 °C with primary antibodies rabbit anti-YAP1 (LS Bio, cat # LS-C331201-20) at a dilution of 1/300 in diluted blocking buffer (dBB, 1/5 dilution of blocking buffer in PBS-T). The following day, cells were washed with PBS-T and incubated for 35 min at RT with secondary antibodies A647 goat anti-rabbit (Invitrogen, Cat # A21245) and A546 goat anti-rat (Invitrogen, Cat # A110819) at a dilution of 1/500 in dBB. Secondary antibodies were later washed with PBS-T and PBS and after that, cells were incubated for 5 minutes at RT with DAPI (5 μg/ml) followed by washing steps with PBS-T and PBS. Cells were imaged in PBS using a Zeiss LSM780 confocal microscope using 40X objective.

### Phagocytosis assay

For the phagocytosis assay, cells were cultured in 96-well plate and treated with conjugates as previously described (for 4 h at a 10 μM concentration at 37°C, then peptide conjugates were removed, washed with medium and fresh medium was added). The phagocytosis assay was performed at 48 h after treatment removal by removing the medium, incubating fluorescent E. coli BioParticles for 2 h, followed by a 1 min incubation with trypan blue (Vybrant™ Phagocytosis Assay Kit, Invitrogen, Cat# V6694). Plates were read at 480 nm excitation and 520 nm emission using a BioTek Synergy H1 plate reader.

### MACTIDE-V monotherapy in orthotopic 4T1.2

5x10^5^ 4T1.2 cells in 50 µL of PBS was injected subcutaneously to 4^th^ mammary fat pad. On day 7, mice were sorted into groups based on tumor volume, approximately 55-60 mm^3^ calculated based on the formula shown above. Mice were treated every other day with 500 µL of MACTIDE-V, CtrlPep-V (30 nmoles) and PBS, 9 injections in total, *n*=10 mice/group. Cumulative Verteporfin dose was 9 mg/kg. The last injection was on day 23. Four mice from each group were sacrificed on day 25 through anesthetic overdose and their tumors analyzed by flow cytometry (FC) and mIHC. The rest of the mice were left to continue in the study with the initial intention of performing a survival study, however, all of these mice were sacrificed within five days based on their tumor size (above 1500 mm^3^) or cachexia.

### Flow cytometry

For FC, mice were not perfused, the tumors were cut into small pieces, digested using 10 mL of collagenase IV (200 U/mL, Gibco™, catalog no.17104019), dispase (0.6 U/mL, Gibco™, catalog no. 17105-041) and DNase I (15 U/mL, AppliChem, catalog no. A3778) mixture on a rotating platform for up to 60 min at 37 °C, pipetting every 10 min. Red blood cells were lysed using 3 mL of ammonium-chloride-potassium lysis buffer. After that, cells were centrifuged (350 x g, 7 min, +4 °C), filtered (100 µm cell strainer, Falcon, catalog no. 352360) and counted using the bright-field mode of LUNA Automated Cell counter (Logos Biosystems). Cells were seeded at a concentration of 5 x 10^6^ cells/100 µL of running buffer (RB) (1L of RB: 4 ml 0.5M EDTA + 100ml FBS + the rest PBS) on 96-well conical bottom plate and incubated for 15 min in dark at RT in 50 µL of Zombie NIR™ Fixable Viability Kit (Biolegend). After that, 30 µL of blocking antibody (TruStain FcX, Biolegend) was added and incubated 10 min in dark at +4 °C. Then, to stain macrophages, 20 µL of antibody mixture was added, incubated 25 min in dark at +4 °C after which 50 µL of RB was added, centrifuged, washed two times with 150 µL of RB and taken up in 150 µL of RB. For intracellular staining, cells were fixed and permeabilized using the eBioscience™ Foxp3 / Transcription Factor Staining Buffer Set (Thermo). Cells were then stained with 100 µL of antibody mixture and incubated 40 min in dark at RT. Cells were centrifuged, washed twice with 150 µL of RB and resuspended in 150 µL of RB. Samples were analyzed using the SONY ID7000 spectral flow cytometer and the accompanying software.

### Antibodies used for *in vivo* FC

| Antigen | Fluorophore | Clone | Catalog number | Company |
| --- | --- | --- | --- | --- |
| CD86 | BV421 | GL-1 | 105032 | Biolegend |
| PD-1 | BV510 | 29F.1A12 | 135241 | Biolegend |
| LY6C | SBV570 | ER-MP20 | MCA2389SBV570 | Bio-Rad |
| CD11b | BV750 | M1/70 | 101267 | Biolegend |
| CD3 | FITC | 145-2C11 | 100306 | Biolegend |
| MHC II | NovaFluorBlue610 | M5/114.15.2 | # M024T02B06 | Thermofisher |
| F4/80 | SBB765 | Cl:A3-1 | MCA497SBB765 | Bio-Rad |
| CD45 | SBB810 | YW62.3 | MCA1031SBB810 | Bio-Rad |
| CD206 | PE-Fire700 | C068C2 | 141742 | Biolegend |
| PDL1 | NovaFluorRed700 | MIH5 | M036T03R03 | Thermofisher |
| CD11b | APC-Cy7 | N418 | 117324 | Biolegend |
| B220 | APC-Fire810 | RA3-6B2 | 103278 | Biolegend |
| GRZB | BV421 | QA18A28 | 396414 | Biolegend |
| CD49 | PacBlue | DX5 | 108918 | Biolegend |
| CD4 | BV570 | RM4-5 | 100542 | Biolegend |
| CD27 | BV605 | LG 3A10 | 124249 | Biolegend |
| CD8 | BV650 | 53-6.7 | 100742 | Biolegend |
| CD44 | BV711 | IM7 | 103057 | Biolegend |
| FOXP3 | AF488 | 150D | 320012 | Biolegend |
| CD3 | SparkBlue-574 | 17A2 | 100276 | Biolegend |
| CD19 | SBB765 | 6D5 | MCA1439SBB765 | Bio-Rad |
| IL7RA | PE-Cy7 | A7R34 | 135014 | Biolegend |
| CD25 | APC | PC61 | 102012 | Biolegend |
| KI67 | AF700 | 16A8 | 652420 | Biolegend |
| CD62L | APC-Cy7 | MEL-14 | 104428 | Biolegend |

For FC of mouse BMDMs, macrophages were detached at day 7, 48 h after treatment with either PBS or PDCs, with cold PBS 2mM EDTA. Cells were pelleted, washed and stained in PBS 20′ at 4°C with the LIVE/DEAD™ Fixable Scarlet (723) Viability Kit (Cat# L34986, Invitrogen, ThermoFisher Scientific) and the antibodies listed below.

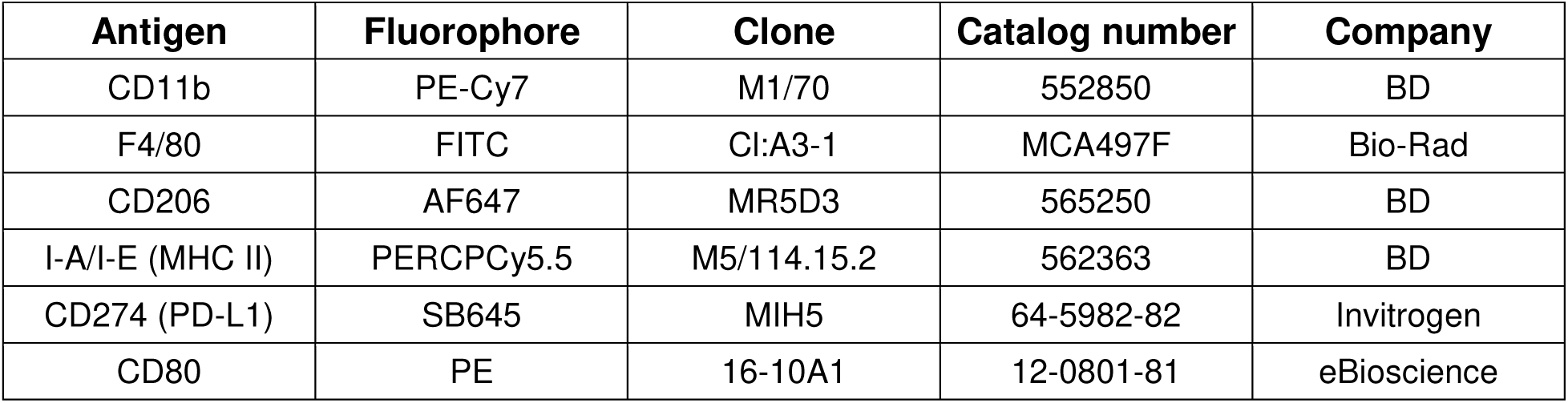

### Metastasis assessment

To determine the pulmonary metastatic area, lungs from MACTIDE-V monotherapy were stained for hematoxylin and eosin according to the following protocol. Unfixed slides (10 µm in thickness) were taken out of the -20 °C freezer 10 min before starting the staining. First, slides were fixed 2 min in ice cold methanol, then incubated 3 min in hematoxylin solution after which slides were washed 5 min with running tap water. Then, slides were incubated for 3 min in eosin solution after which washing 5 min in running water followed. For rehydration, slides were placed into two times 100% ethanol for 1 min and then for clearance two times into RotiClear (Roth, catalog no. A538.5) for 2 min after which slides were mounted using Eukitt quick hardening mounting medium (Merck, catalog no. 03989). Slides were then scanned using Leica DM6 B microscope and Leica Aperio Versa 8 slide scanner with 20x zoom. Images were analyzed using the QuPath program to determine the pulmonary tumor area coverage by dividing the tumor area per whole lung area and multiplying with 100.

### MACTIDE-V + anti-PD1 combination therapy in orthotopic 4T1.2

5x10^5^ 4T1.2 cells in 50 µL of PBS were injected subcutaneously into 4^th^ mammary fat pad. On day 8, mice were sorted into groups based on tumor volume (approximately 40 mm^3^): PBS, anti-PD-1, MACTIDE-V, MACTIDE-V+anti-PD-1. Mice were treated i.p. every other day with 500 µL of MACTIDE-V (30 nmoles, 1mg/kg Verteporfin) or PBS (10 injections in total). On day 18 post injection, MACTIDE-V+anti-PD-1 and anti-PD-1 groups received recombiMAb anti-mouse PD-1 (Bioxcell, Catalog #CP151), a mouse IgG2a monoclonal antibody with the D265A mutation in the Fc fragment which renders it unable to bind to endogenous Fcγ receptors. Mice received three intraperitoneal injections of 200 µg of anti-PD-1 dissolved in 500 µL of PBS. The groups that did not receive anti-PD1 were injected with 500 µL of PBS i.p. The last injection was on day 27, all mice were sacrificed on day 29 through anesthetic overdose. Five tumors per group were analyzed by FC as described above. TAM/monocytes were defined as Ly6G^-^CD11b^+^/F4/80^+^ population from the CD45^+^ population.

The lungs of all mice were fixed in formalin and embedded in paraffin, and later sectioned and stained with H&E by the histology facility of Vall D’Hebron Research Institute (Barcelona, Spain). Five 10 μm-sections spaced apart 1 mm were taken from the lungs of each mouse (*n*=7 mice per group). Then, the slides were scanned, and the metastatic area divided by the total lung area was calculated for each mouse.

### Data Availability

All data needed to evaluate the conclusions on the article are presented in the article and/or the Supplementary Data. Additional data related to the findings of this study are available from the corresponding author.

## Acknowledgements

P.S. acknowledges support from MICIN/AEIMCIN/AEI/10.13039/501100011033/ and by “ERDF A way of making Europe”, (grant PID2021-122364OA-I00; and Ramon y Cajal contract RYC2020-028754-I), the Estonian Research Council (grant PUT PSG38) and the Royal Society of Chemistry (Research Fund Grant R21-6405053838). I.M. acknowledges Università di Padova (grant DiSCOG MARI_BIRD22_01). M.R. acknowledges support from CIBER BBN(CB06/01/0074). P.P. acknowledges support from the Estonian Research Agency (grant PRG2011). T.T. was funded by the Estonian Research Council (grants PRG230 and PRG1788), MSCA-RISE 2018 Project OXIGENATED, and EuronanomedIII (projects ECM-CART and iNanoGun).

## Author contributions

A.L. performed in vitro and in vivo experiments, designed experiments, wrote the manuscript, provided analysis and discussion. E.P. performed in vitro experiments with BMDMs, designed experiments, wrote the manuscript and provided discussion and analysis. U.H. performed flow cytometry studies from in vivo experiments, designed experiments, edited the manuscript and provided analysis and discussions. E.K.A. performed modeling studies. M.C.A. and L.M. designed, analyzed and performed in vitro experiments with BMDMs. K.K. performed LC-MS studies. M.L, M.R, G.A. provided peptide and peptide-conjugate synthesis. I.M. and P.P. provided funding and discussion. T.T. provided funding and discussion and edited the manuscript. P.S. performed binding and homing studies, designed experiments and provided supervision, provided funding and wrote the manuscript.

