## Supporting information for "Therapeutic Tumor Macrophage Reprogramming in Breast Cancer Through a Peptide-Drug Conjugate"

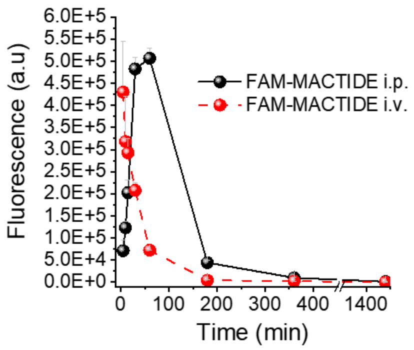


**Figure S1.**: Pharmacokinetics of i.p. and i.v. administered FAM-MACTIDE (30 nmoles) analyzed by detecting plasma fluorescence in the FAM channel at different timepoints (N=3).

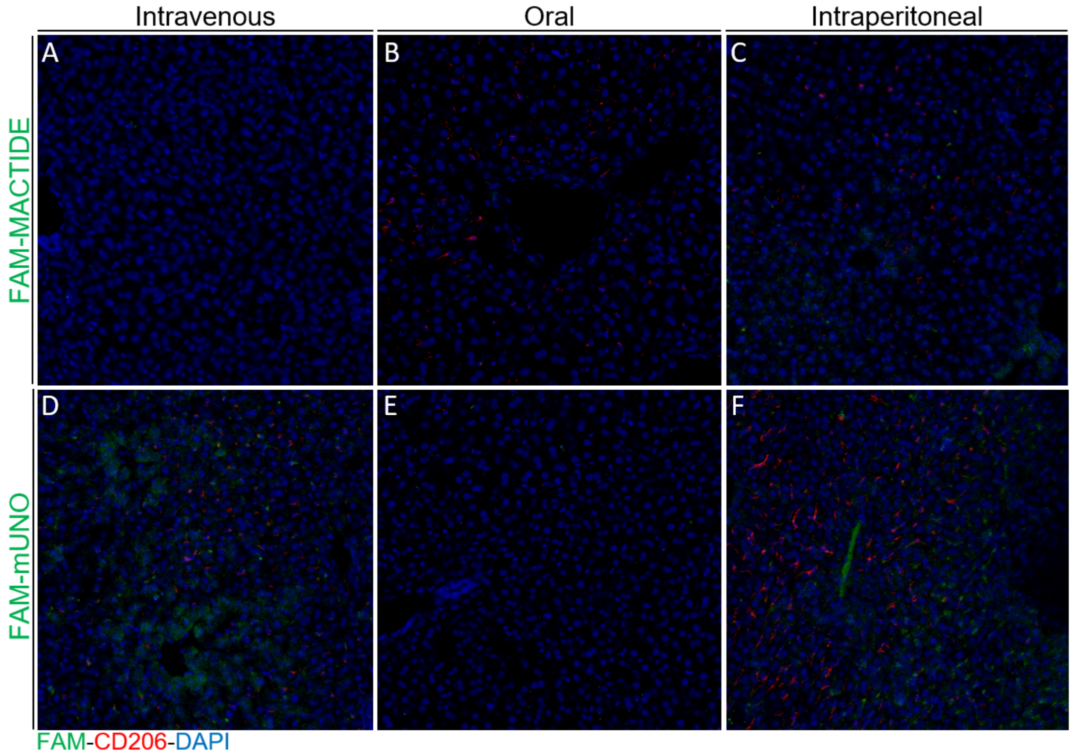


**Figure S2**. Intravenously, intraperitoneally and orally administered FAM-MACTIDE or FAM-mUNO does not accumulate in the liver of breast cancer in mice. 30 nmoles of FAM-MACTIDE or FAM-mUNO were administered and left to circulate for 24 h. At 24 h, the mice were sacrificed, and the organs were collected, fixed, cryoprotected, sectioned, and immunostained for FAM (shown in green) and CD206 (shown in red). Representative images from n=3 mice are shown.

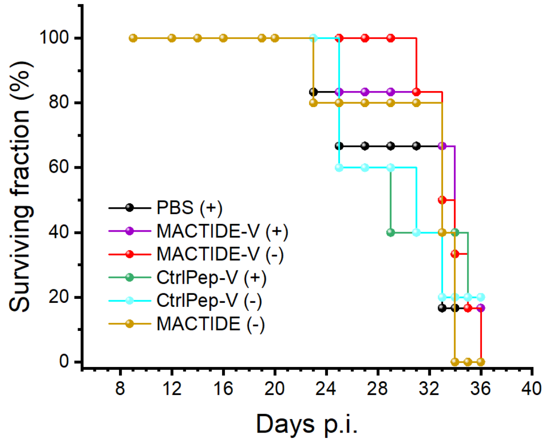


**Figure S3**. Survival for treatment study of figure 5.

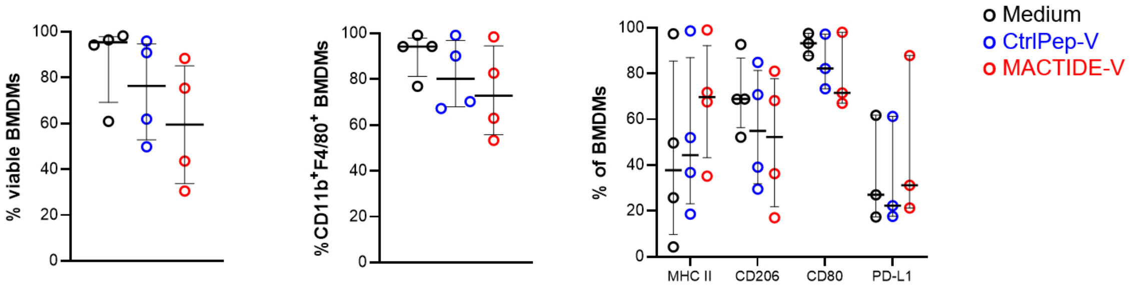


**Figure S4. Flow cytometry on BMDMs 48 hours after treatment**. *Left panel*, percentage of viable cells in Balb/c BMDM cultures. *Central panel*, percentage of CD11b$^+$F4/80$^+$ BMDMs in BALB/c cultures. *Right panel*, percentage of MHC II$^+$, CD206$^+$, CD80$^+$ and PD-L1$^+$ BMDMs in BALB/c samples. Median ± interquartile range. Repeated measures one-way ANOVA with multiple comparisons for *n*=4 independent experiments.

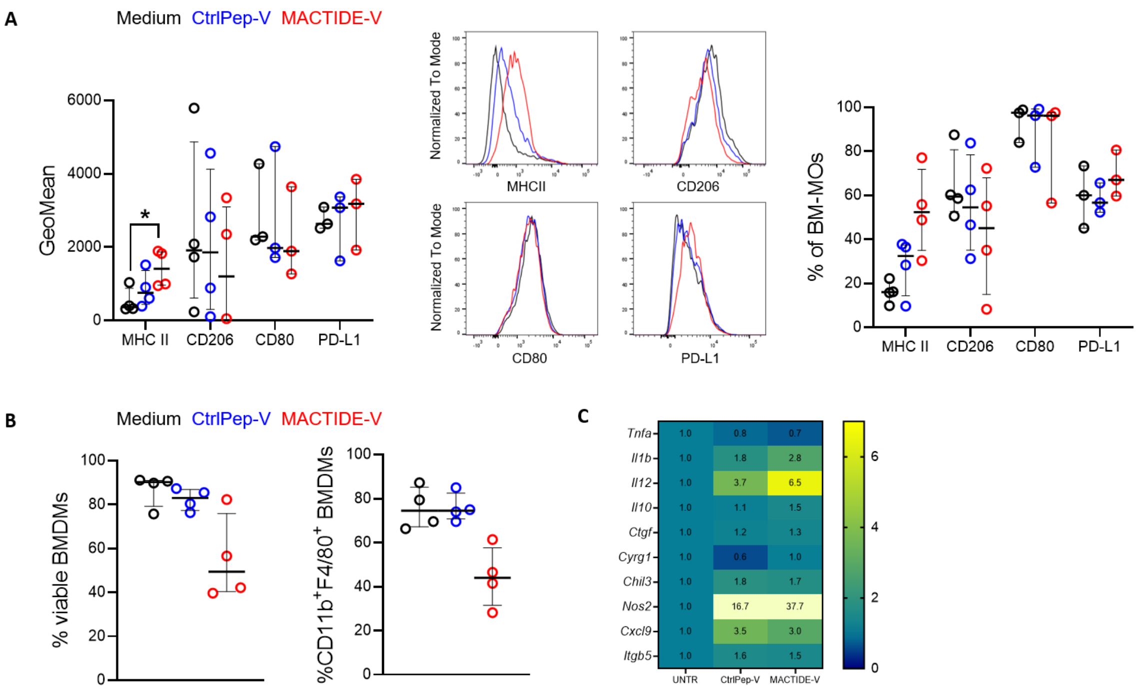


**Figure S5. (A)** Flow cytometry on BMDMs 48 hours after treatment. *Left panel*, GeoMean of MHC II in B6 BMDMs. Median ± interquartile range. Repeated measures one-way ANOVA with multiple comparisons, n=4 independent experiments. *$p \leq 0.05$. *Central panel,* concatenated GeoMean histograms for MHC II in B6 BMDMs, *n*=4. *Right panel,* percentage of MHC II$^+$, CD206$^+$, CD80$^+$ and PD-L1$^+$ BMDMs in B6 samples. Median ± interquartile range. Repeated measures one-way ANOVA with multiple comparisons, *n*=4 independent experiments. **(B)** *Left panel,* percentage of viable cells in B6 BMDM cultures. *Right panel*, percentage of CD11b$^+$F4/80$^+$ BMDMs in B6 cultures. Median ± interquartile range. Repeated measures one-way ANOVA with multiple comparisons, n=4 independent experiments. **(C)** Heatmap of the mRNA expression of genes involved in the functional activation of BMDMs, YAP signaling and adhesion, measured by real-time PCR 48 hours after treatment in B6 BMDMs, *n*=2 independent experiments.

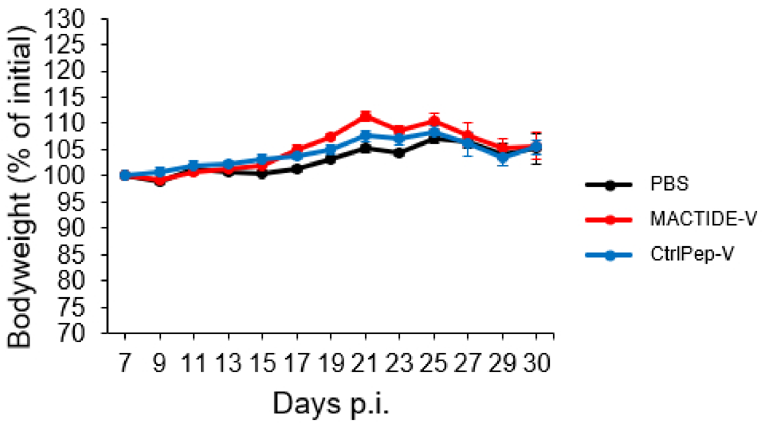


**Figure S6.** Bodyweights for treatment study of figure 7.

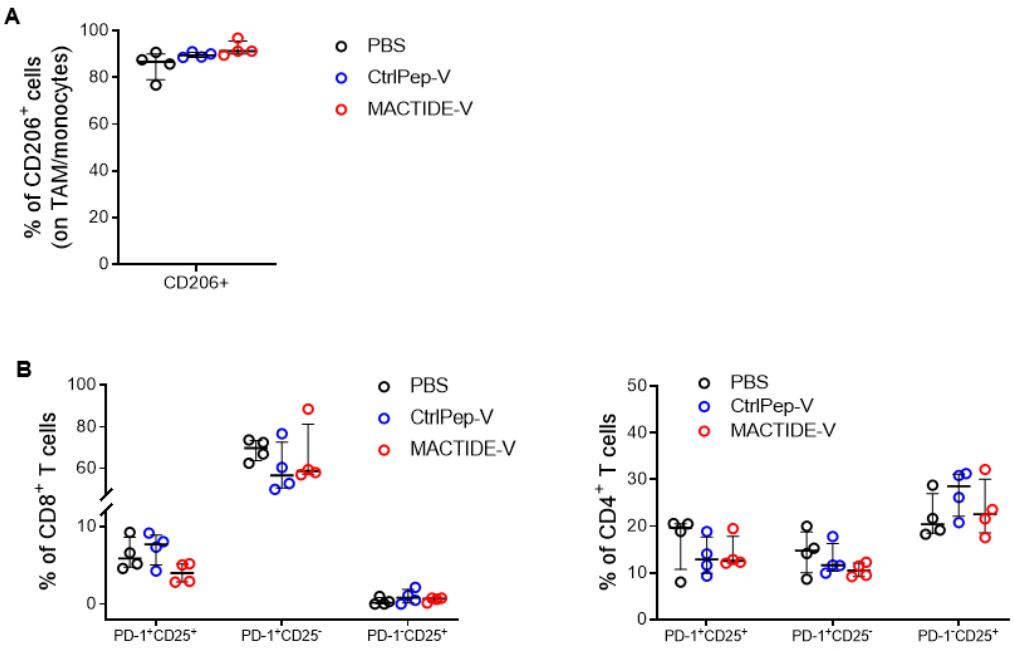


**Figure S7.** Flow cytometry of the orthotopic 4T1.2 tumors of Figure 7, *n*=4.

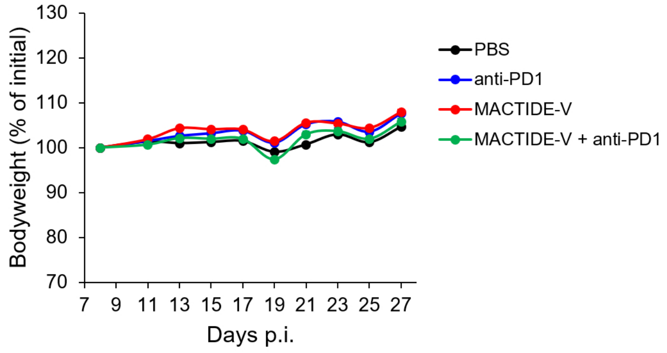


**Figure S8.** Bodyweights for treatment study of figure 8.

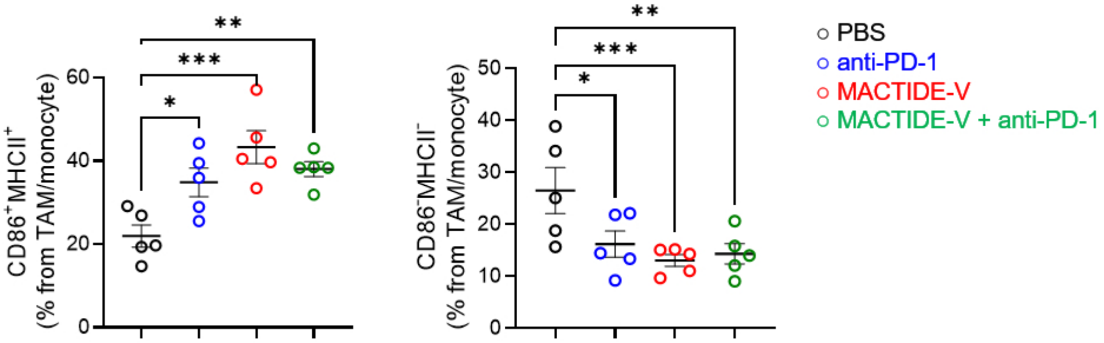


**Figure S9.** Flow cytometry of the orthotopic 4T1.2 tumors of Figure 8, *n*=5.
